# Full structure/function analysis of all the pilin subunits in a type 4 pilus: a complex of minor pilins in *Streptococcus sanguinis* mediates binding to glycans

**DOI:** 10.1101/2022.08.25.505150

**Authors:** Meriam Shahin, Devon Sheppard, Claire Raynaud, Jamie-Lee Berry, Ishwori Gurung, Lisete M. Silva, Ten Feizi, Yan Liu, Vladimir Pelicic

## Abstract

Type 4 filaments (T4F) – of which type 4 pili (T4P) are the archetype – are a superfamily of filamentous nanomachines nearly ubiquitous in prokaryotes. T4F are polymers of one major pilin that also contain minor pilins whose roles are often poorly understood. Here, we complete the structure/function analysis of the full set of T4P pilins in the opportunistic pathogen *Streptococcus sanguinis*. We determined the structure of the minor pilin PilA, which is unexpectedly similar to one of the subunits of a tip-located complex of four minor pilins, widely conserved in T4F. We found that PilA interacts and dramatically stabilises the minor pilin PilC. We determined the structure of PilC, showing that it is a modular pilin with a lectin module binding a specific subset of glycans prevalent in the human glycome, the host of *S. sanguinis*. Altogether, our findings support a model whereby the minor pilins in *S. sanguinis* T4P form a tip-located complex promoting adhesion to various host receptors. Our findings have general implications for a group of minor pilins widely conserved in T4F.

## Introduction

Type 4 filaments (T4F) – type 4 pili (T4P) and type 2 secretion systems (T2SS) being the best known – are a superfamily of filamentous nanomachines ubiquitous in prokaryotes^1,2^. T4F, which mediate a wide variety of functions^1^, are composed of type 4 pilins and are assembled by conserved multi-protein machineries spanning the cell envelope^3^. T4F have been studied for decades because they are virulence factors in many human pathogens.

Our current understanding of T4F is as follows. T4F are polymeric filaments of type 4 pilins, one predominant (major) and several low abundance (minor) subunits^4^. Pilins are synthesised as prepilins with a distinctive N-terminal sequence motif – known as class 3 signal peptide (SP)^5^ – consisting of a short hydrophilic leader peptide ending with a Gly or Ala, followed by a hydrophobic tract of 21 residues. Pilins display a characteristic “lollipop” structure where the hydrophobic tract of the SP constitutes the N-terminal portion (α1N) of an α-helix of approx. 50 residues (α1) that supports a globular head^4^. However, larger minor pilins – displaying extra domains “grafted” onto a pilin moiety and hence defined as modular pilins^6^ – are not uncommon, which partly explains T4F extreme functional versatility. Pilins are assembled in filaments at the cytoplasmic membrane^3^ in two steps. The leader peptide in prepilins is first cleaved by a dedicated prepilin peptidase (PPase)^7–9^, before the pilins are polymerised by a machinery centred on a platform protein and a hexameric ATPase^10^. All T4F are helical polymers where pilins are held together by extensive interactions between their α1-helices, which pack within the core of the filament^11^. However, slightly different helical symmetry parameters – axial rise and azimuthal rotation between consecutive subunits – are seen in different T4F^12–16^.

While the role of major pilins is clear, *i.e.*, they constitute the backbone of T4F, the role and precise localisation of the minor pilins remain poorly defined^4^. Typically, minor pilins are divided into two categories: widely conserved and non-conserved (or system-specific)^4^. Non-conserved minor pilins are highly variable, and their roles cannot be predicted. In some T4F, these pilins have been well studied, such as for example ComP, PilV, PilX in *Neisseria meningitidis* T4P. These three pilins are dispensable for filament assembly but are key for different T4P-mediated functions. ComP binds DNA to promote its uptake during natural transformation^17,18^, PilV promotes bacterial adhesion to human cells^19^, and PilX supports the formation of bacterial aggregates^20^. In contrast, a group of four minor pilins – often labelled with the letters H, I, J, K – are widely conserved in different T4F and essential for filament assembly^21–23^. Their role and mechanism of action are likely to be the same since HIJK sets from different T4F are functionally interchangeable in promoting filament assembly^24^. These pilins interact to form a quasihelical complex^25,26^, with symmetry parameters consistent with T4F. This complex is capped by the bulky K subunit, suggesting a localisation at the pilus tip^25,26^, which was recently confirmed by cryo-electron tomography^27^ (cryo-ET) in *Myxococcus xanthus* T4P. Since T4F are assembled from tip to base^3^, the HIJK complex is therefore thought to initiate filament assembly^28,29^.

Recently, the study of T4F in monoderm bacteria opened new research avenues^30^, with *Streptococcus sanguinis* becoming a cutting edge model^31^. This commensal of the human oral cavity^32^ is an opportunistic pathogen frequently causing infective endocarditis (IE)^33^. It uses an elementary machinery (with fewer components) to assemble T4P composed of two major (PilE1, PilE2) and three minor (PilA, PilB, PilC) pilins^34,35^. This apparent simplicity makes *S. sanguinis* T4P an ideal system to characterise the role of all pilin subunits in detail, which is yet to be achieved in any T4F. Since we previously characterised PilE1, PilE2 and PilB^6,34,35^, here we focused on the minor pilins PilA and PilC. We report the detailed structure/function analysis of these two proteins, how they interact and how they function. We provide the first integrated view of the role of all the pilin subunits in a defined T4F, which has implications for this widespread superfamily of filamentous nanomachines.

## Results

### PilA and PilC are minor pilins with remarkably different characteristics

Previously, we showed that all the genes involved in T4P biology in *S. sanguinis* 2908 strain cluster together in a 22 kb locus named *pil*^34^. This locus encodes five pilins – PilE1, PilE2, PilA, PilB, PilC (Fig. 1A) – which are essential for piliation^34^. These proteins are processed by the PPase PilD and become subunits of T4P^34,35^. PilE1 and PilE2 are the major pilins, whereas PilA, PilB and PilC are minor pilins^35^. We previously characterised PilE1, PilE2 and PilB^6,35^. In this study, we complete an exhaustive analysis of the role of pilins in T4P biology in *S. sanguinis* 2908 by focusing on PilA and PilC.

**Fig. 1.**
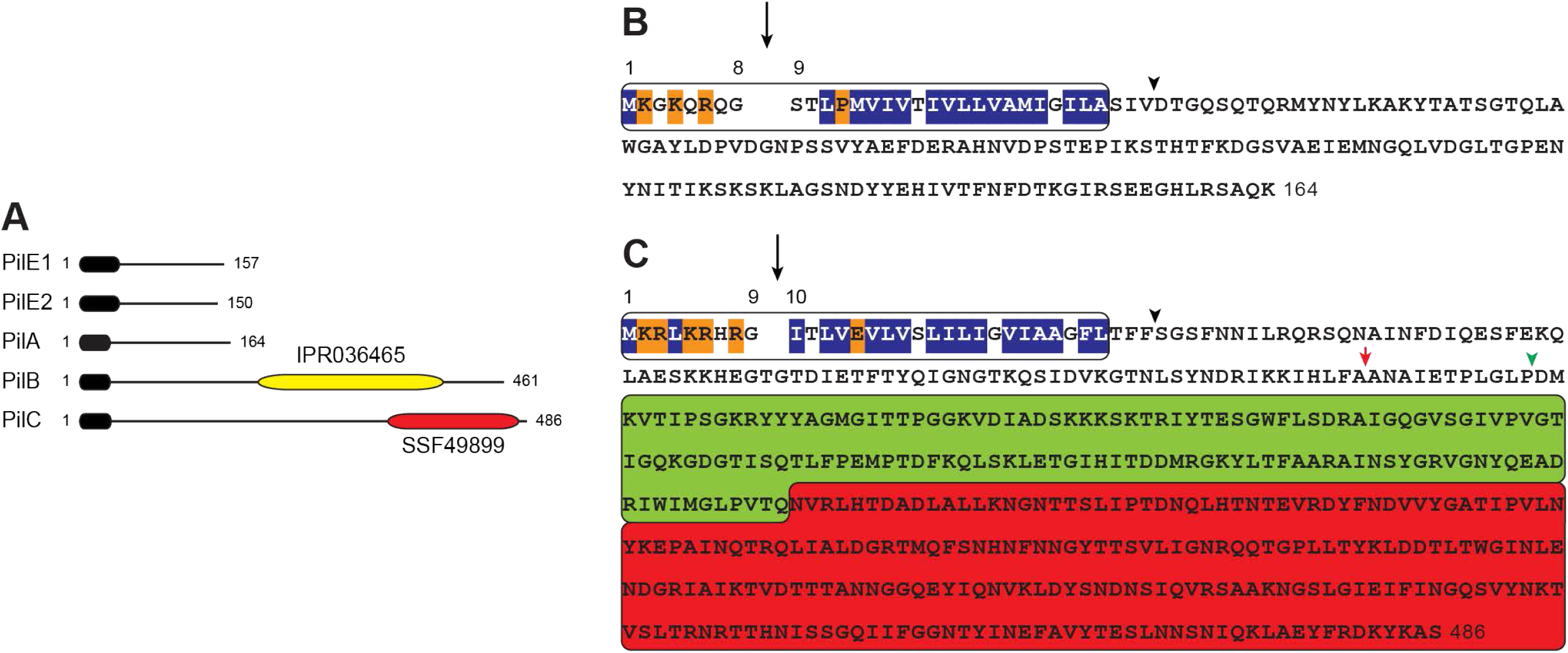
PilA and PilC sequence characteristics. **A**) Protein architecture of the five pilins in the T4P-encoding *pil* locus in *S. sanguinis* 2908^34^. PilE1 and PilE2 are the major pilins, while PilA, PilB, and PilC are minor pilins. Each protein harbours a type 4 pilin-defining class 3 SP (black rounded rectangle) at its N-terminus. PilB and PilC are modular pilins, which contain additional domains at their C-terminus (yellow and red rounded rectangles). PilB contains a von Willebrand factor A-like domain (IPR036465)^6^, while PilC contains a concanavalin A-like lectin/glucanase structural domain (SSF49899). Proteins are drawn to scale. **B**) Relevant sequence features of PilA. The class 3 SP is boxed and contains an 8-aa long leader peptide with mostly hydrophilic (orange shading) and neutral (no shading) residues, followed by a tract of 21 mainly hydrophobic residues (blue shading). The processing of the leader peptide by the PPase, indicated by the vertical arrow, generates a pilin of 156 residues (17 kDa). The arrowhead indicates the portion of PilA that was produced and purified in this study. **C**) Relevant sequence features of PilC. The class 3 SP is boxed. The processing of the leader peptide by the PPase (black vertical arrow), generates a 477-aa long pilin (52.8 kDa). The lectin module (SSF49899) is boxed and coloured in red. The Ig-like fold module revealed by our structure is boxed and coloured in green. Arrowheads indicate the proteins that were produced and purified in this study, with/without the pilin moiety (black and green arrowheads, respectively). The red vertical arrow indicates the site of spontaneous proteolysis of purified 6His-PilC.

Although it has a typical pilin size – 17 kDa for the processed protein – PilA is an unusual pilin. Its class 3 SP, which was identified by visual inspection^34^, is degenerate (Fig. 1B) and could not be identified by any of the available bioinformatic tools, including PilFind dedicated to this purpose^36^. This is mainly due to the presence of a Pro residue in the fourth position of the processed protein (Fig. 1B), which is very uncommon. No prediction could be made about the role of PilA, as this protein has no detectable sequence homologs outside of *S. sanguinis*.

PilC has radically different features. It has a canonical class 3 SP (Fig. 1C), with a typical Glu residue in the fifth position of the processed protein^23^. Processed PilC is unusually large for a pilin, with a theoretical molecular mass of 52.8 kDa. As recently reported for PilB^6^, the large size of PilC is due to its predicted modular architecture, with a well-defined domain not specific to T4P biology grafted onto the C-terminus of a pilin moiety (Fig. 1C). Whereas in PilB the extra module corresponds to a vWA domain (IPR036465)^6^, in PilC it belongs to the concanavalin A-like lectin/glucanase domain superfamily (SSF49899). This superfamily is composed of very diverse proteins in the three domains of life, ranging from legume lectins (*e.g.*, concanavalin A) to animal lectins (*e.g.*, galectins). All these proteins are carbohydrate-binding proteins, but they differ in their ligand specificities. These observations raise the possibility that PilC is a modular pilin with a lectin module promoting T4P-mediated adhesion of *S. sanguinis* to glycans.

### PilA is structurally similar to a widely conserved minor pilin from a complex of minor pilins found at the tip of many T4F

To better understand the role of PilA, we solved its 3D structure by X-ray crystallography. We produced a 15.7 kDa recombinant protein in *Escherichia coli*, in which the N-terminal 32 residues of PilA (encompassing the leader peptide and hydrophobic α1N) (Fig. 1B) were replaced by a hexahistidine tag (6His). α1N truncation promotes pilin solubility without having a structural impact on the rest of the protein^37^. We purified soluble 6His-PilA using a combination of affinity and size-exclusion chromatography (SEC). The protein crystallised readily, and after optimising the best diffracting crystals, we collected a complete dataset in the space group *P*4_3_2_1_2 (Table 1). After phase determination, using crystals produced in the presence of seleno-methionine (SeMet), we solved a 1.77 Å resolution structure. As can be seen in Fig. 2A, PilA exhibits a typical pilin architecture, with a globular head consisting of a long N-terminal α1-helix packed against a β-sheet composed of five anti-parallel β-strands. Modelling the full-length PilA protein using AlphaFold^38^ reveals a canonical lollipop shape^4^ where the globular head is mounted onto a long α1-helix “stick” (Fig. S1). PilA looks like an “ice axe” (Fig. 2A) because of the short α2-helix between α1 and β1, which is almost perpendicular to α1. The almost perfect superposition of our crystal structure with the AlphaFold prediction (Fig. S1) – with a mere 0.58 Å root-mean-square deviation (RMSD) – validates the accuracy of both AlphaFold^38^ and our structure.

**Fig. 2.**
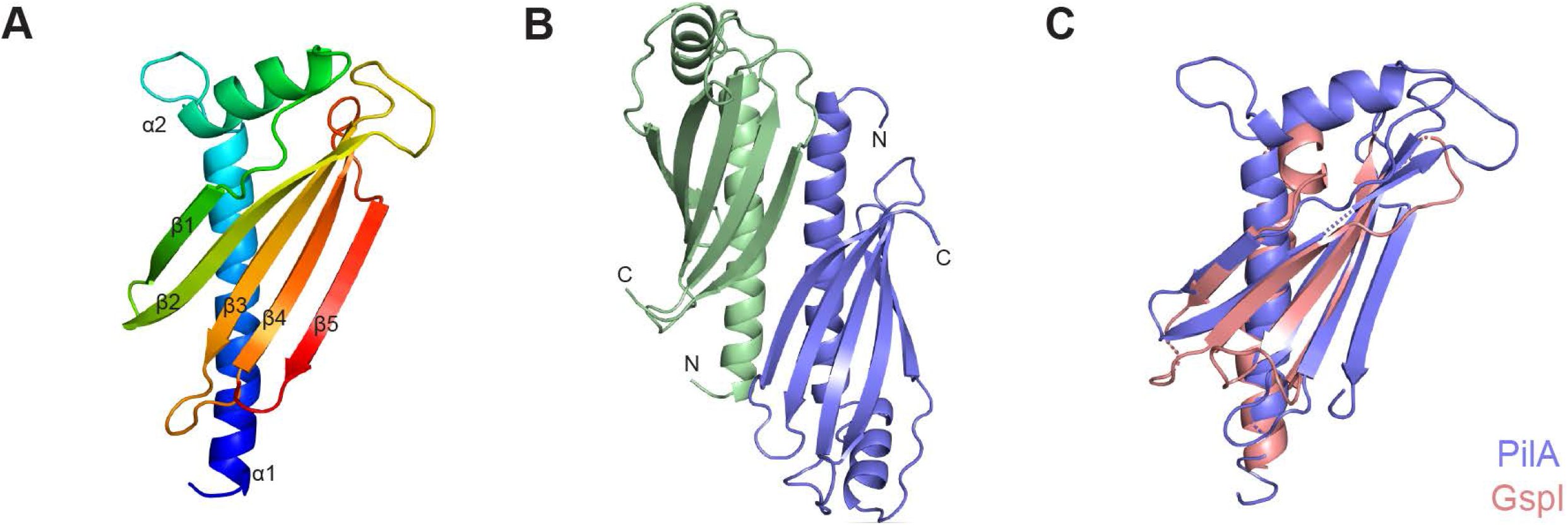
Crystal structure of PilA. **A**) Cartoon view of the PilA structure rainbow-coloured from blue (N-terminus) to red (C-terminus). This highlights a canonical pilin fold with a long N-terminal α-helix (α1) packed against a β-sheet, consisting of five anti-parallel β-strands (β1 to β5). **B**) Cartoon view of the head-to-toe PilA dimers in the crystal packing, which are stabilised by a series of hydrogen bonds between residues in the α1-helices and the last 2 strands of the β-sheets (β4 and β5) (see Fig. S2). This explains why recombinant 6His-PilA purifies as a dimer. **C**) Structural similarity between PilA (blue) and a widely conserved minor pilin in T4F (pink). The GspI protein from enterotoxigenic *E. coli* T2SS (PDB 3CI0) is presented. Although these two proteins share only 9.4 % sequence identity, they superpose with an RMSD of 1.52 Å.

**Table 1.**
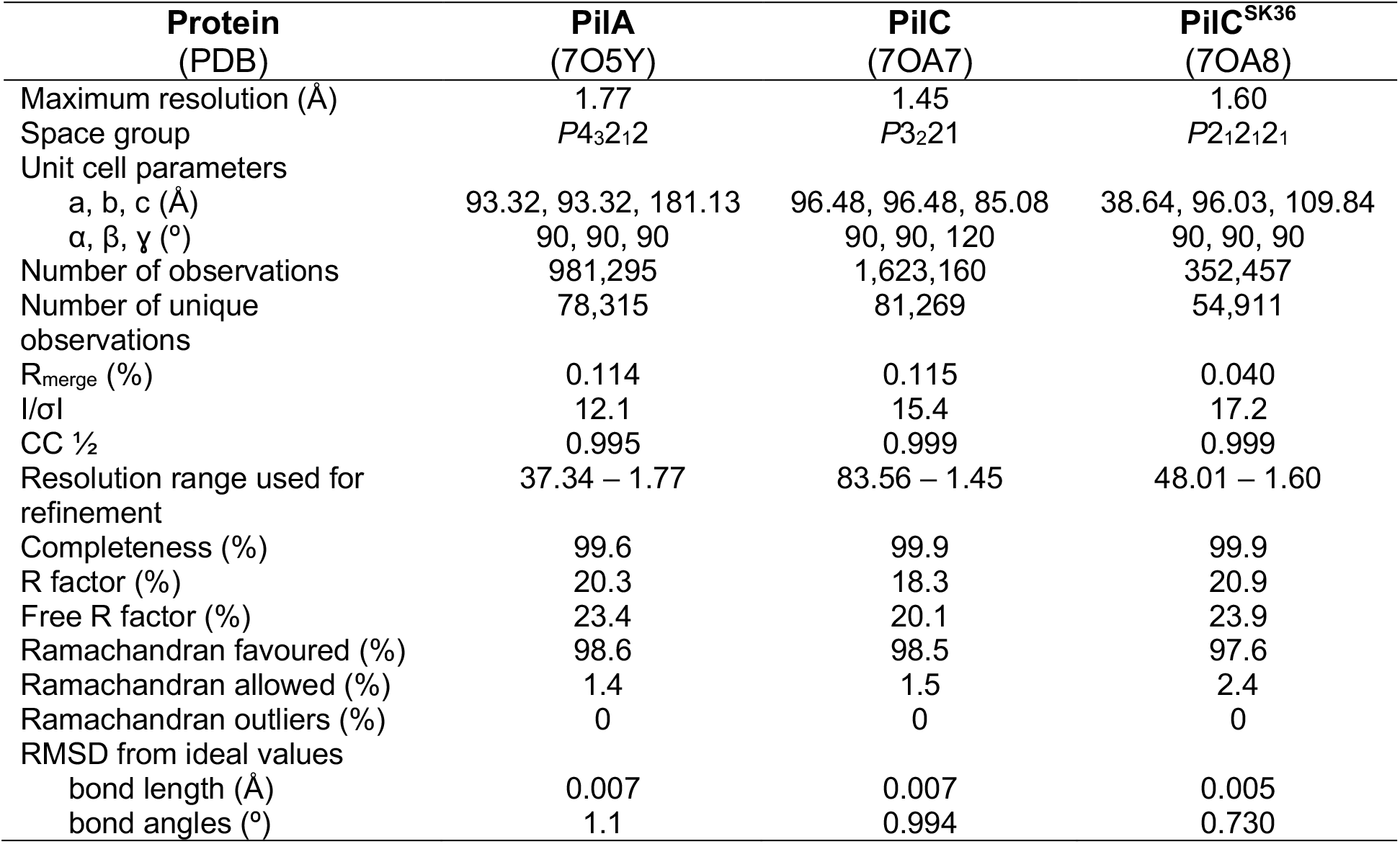
Crystal structures data collection and refinement statistics.

A closer inspection of the crystal packing showed that PilA forms head-to-toe dimers (Fig. 2B), which are stabilised by a series of hydrogen bonds between residues in α1-helix and the last two β-strands of the β-sheet (Fig. S2). This explains why recombinant 6His-PilA purifies as a dimer. Strikingly, while PilA exhibits no sequence homology to other pilins, our structure reveals extensive structural similarity to GspI (PDB 3CI0) from enterotoxigenic *E. coli* T2SS (Fig. 2C). PilA and GspI superpose with an RMSD of 1.52 Å. GspI is part of a HIJK tip-located complex of conserved minor pilins^25,26^, which is widespread in T4F^27^. Critically, GspI interacts with the large GspK^25,26^, which is likely to be a modular pilin although the function of its extra domain is unknown. To us, this similarity suggested that PilA might be interacting with other minor pilins (PilB and/or PilC) to form a complex capping *S. sanguinis* T4P.

### PilA interacts with PilC

To identify interactions between PilA and other pilins in *S. sanguinis* T4P, we performed pull-down assays with proteins purified using two different affinity tags: 6His and Strep-tag II (Trp-Ser-His-Pro-Gln-Phe-Glu-Lys). In this assay, (1) purified proteins corresponding to the soluble portion of the pilins were mixed in solution, (2) the 6His-tagged bait was selectively captured using cobalt-based magnetic beads, and (3) pull-down of the Strep-tagged prey was assessed by immunoblotting using specific anti-Pil antibodies. We verified beforehand that Strep-tagged proteins were not captured using these beads (although PilB exhibited very limited binding). Interestingly, while the 6His-tagged bait proteins were always captured, they pulled down Strep-tagged prey proteins only when the PilA/PilC pair was tested (Fig. 3A). 6His-PilA and 6His-PilC were both able to pull-down their Strep-tagged preys, Strep-PilC and Strep-PilA, respectively (Fig. 3A). These experiments thus reveal an interaction between the minor pilins PilA and PilC.

**Fig. 3.**
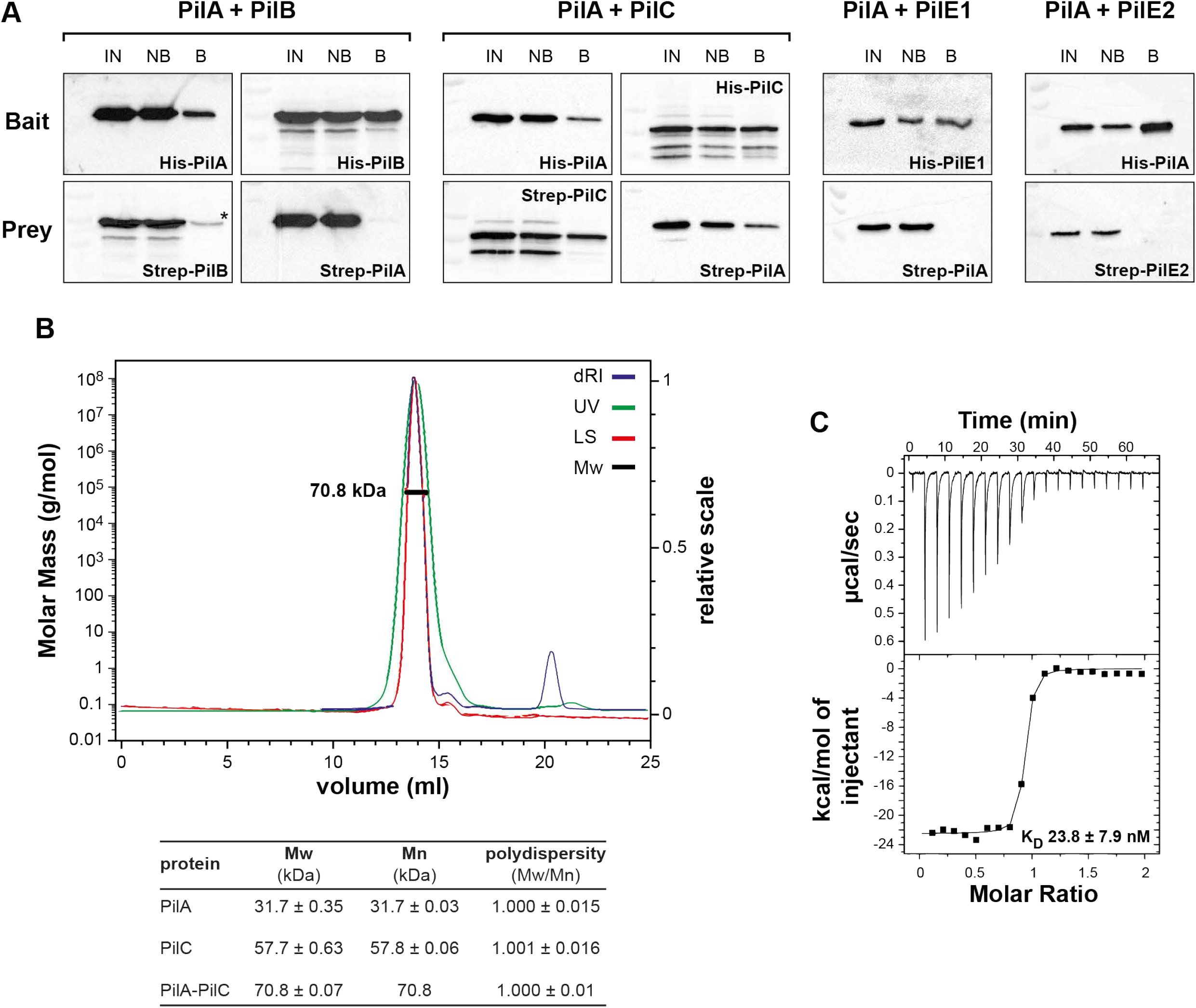
PilA interacts with PilC. **A**) Interactions between PilA and the four other pilins were tested by performing pull-down assays using purified proteins corresponding to the soluble portion of the pilins. The 6His-tagged bait protein was first incubated with Strep-tagged prey protein, and then pulled-down using cobalt-based magnetic beads. Proteins bound to the beads were eluted and identified by immunoblotting using specific anti-Pil antibodies. IN, input. NB, not bound. B, bound. *, non-specific binding of Strep-PilB to the beads, which was also observed with the protein on its own. **B**) Characterisation of the interaction between PilA and PilC by SEC-MALS. Upper panel, chromatogram traces for the PilAC complex. dRI, differential refractive index, LS, MALS signal. Mw, weight average molecular weight. Lower panel, SEC-MALS data for PilA, PilC and PilA-PilC. Mn, number average molecular weight. **C**) Quantification of the PilA-PilC interaction by ITC. A representative ITC titration curve (upper panel) and binding isotherm (lower panel) are presented. The calculated K_D_ is the average ± SD from three independent experiments.

To characterise the PilAC complex, we used SEC coupled with multi-angle light scattering (SEC-MALS). This allows for absolute characterisation of proteins and complexes in solution, in terms of molecular mass and stoichiometry. SEC-MALS was first used with purified 6His-PilA and 6His-PilC on their own. Both proteins eluted as single, monodisperse peaks with estimated molecular masses (in kDa) of 31.7 ± 0.35 and 57.7 ± 0.63, respectively (Fig. 3B). While this value is reasonably close to the theoretical molecular mass for PilC (51.3 kDa), it is much higher for PilA (15.7 kDa). This shows that purified 6His-PilA behaves as a dimer in solution, which probably corresponds to the head-to-toe dimer observed in the crystals (Fig. 2B). When SEC-MALS was performed after mixing equimolar amounts of purified 6His-PilA and 6His-PilC, we observed a single, monodisperse peak with an estimated molecular mass of 70.8 ± 0.07 kDa (Fig. 3B). This value indicates that the PilAC complex consists of one copy of each protein (1:1 stoichiometry). The disappearance of the PilA-PilA homodimer shows that PilC is the preferred binding partner of PilA. Using isothermal titration calorimetry (ITC), we measured the affinity of PilA for PilC (Fig. 3C), which is high, with a dissociation constant (K_D_) of 23.8 ± 7.9 nM.

We then assessed how PilA and PilC interact. We first performed pull-down assays, as above, using a purified 6His-PilC_Δpilin_ protein lacking the pilin moiety (Fig. 1C). In contrast to 6His-PilC (Fig. 3B), the shorter 6His-PilC_Δpilin_ protein was unable to pull-down its Strep-PilA partner (Fig. 4A). This suggests that PilA interacts with the pilin moiety of PilC. Next, we used multidimensional nuclear magnetic resonance (NMR) to identify the portion of PilA binding to PilC. First, we performed a partial NMR assignment of the resonances in 6His-PilA (Fig. S3), highlighting the assigned residues on our crystal structure (Fig. 4B). Then, we identified the chemical shift perturbations occurring in PilA when PilC was added (Fig. S3), which were due to the formation of the PilAC complex and/or the dissociation of the PilA homodimer. Nine chemical shifts perturbations concerned PilA residues not involved in the formation of the PilA dimer (Fig. S2), which are therefore likely to be involved in the PilA-PilC interaction (Fig. 4B). The PilAC interaction interface involves the α1 and α2 helices and the first 2 β-strands (β1 and β2) of PilA (Fig. 4B). The interaction interfaces in the PilAC complex (Fig. 4B) and in the PilA homodimer (Fig. S2) are clearly distinct, since two opposite sides of PilA are involved. We probed the PilAC interaction interface by mutagenesis of residues Ala_76_, Lys_93_ and Thr_95_ in PilA, and quantification by ITC of the ability of these mutant proteins to interact with PilC (Fig. 4C). This showed that PilA_T95A_ displayed a significantly reduced affinity for PilC, corresponding to 38.3 ± 5.9 % of the wild-type (WT) binding.

**Fig. 4.**
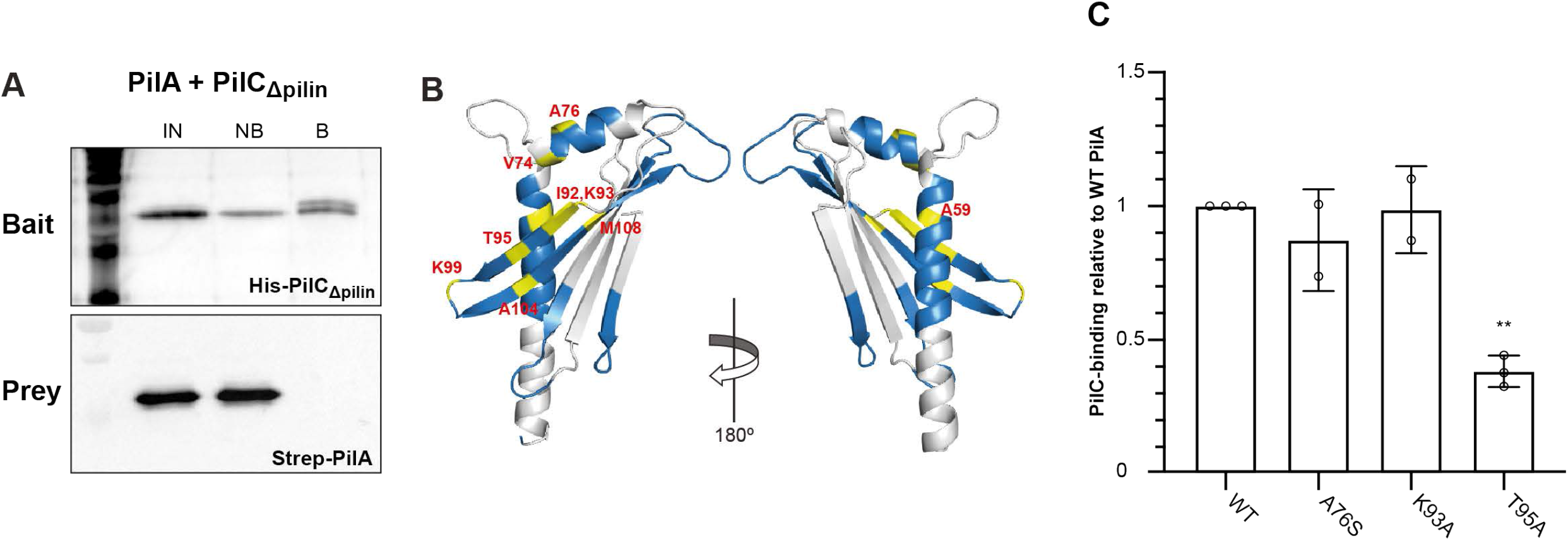
Mapping the PilA-PilC binding interface. **A**) Testing the interaction between PilA and PilC_Δpilin_ by performing pull-down assays as in Fig. 3B. PilC_Δpilin_ corresponds to the protein missing the pilin moiety. Proteins bound to the beads were eluted and identified by immunoblotting using anti-6His and anti-PilA antibodies. IN, input. NB, not bound. B, bound. **B**) NMR analysis of PilAC complex formation. PilA residues for which NMR assignment could be performed are highlighted in blue on our crystal structure (180° view). Residues experiencing significant chemical shift perturbations in the presence of PilC are highlighted in yellow. These residues are in helices α1 and α2, and the first two β-strands. **C**) ITC quantification of the binding of PilA mutant proteins to PilC. The results, presented as K_D_ ratios to the WT, are the average ± SD from 2-3 independent experiments. Statistical significance was calculated using one-way ANOVA followed by Dunnett’s multiple comparison tests.

Taken together, these experiments show that PilA and PilC interact, confirming our hypothesis that PilA is part of a complex of minor pilins in *S. sanguinis* T4P.

### PilA stabilises PilC

We noticed that purified 6His-PilC was unstable, which prompted us to perform long-term stability tests. We kept the purified protein at 4°C, took samples once a week, which we analysed by SDS-PAGE/Coomassie. This revealed that 6His-PilC began degrading during week one and was completely degraded by week two (Fig. 5A), yielding two degradation products of ~ 10 and ~ 40 kDa, respectively. Edman sequencing showed that proteolysis occurred after Ala_110_ (Fig. 1C) close to the end of the pilin moiety, hence the two degradation products were named PilC_Nter_ (smaller N-terminal product) and PilC_Cter_ (larger C-terminal product). While PilC_Cter_ is stable, PilC_Nter_ is highly unstable as it was no longer detectable by week two. In the same conditions, 6His-PilA remained unchanged over four weeks, confirming that is a very stable protein (Fig. 5A). Interestingly, when the PilAC complex was analysed in the same way, PilC degradation was significantly delayed. While PilC on its own was completely degraded by week two, in the presence of PilA, PilC degradation began only at week three, and even at week four the majority of PilC was still intact (Fig. 5A). This was highly reproducible, which indicates that PilA stabilises PilC.

**Fig. 5.**
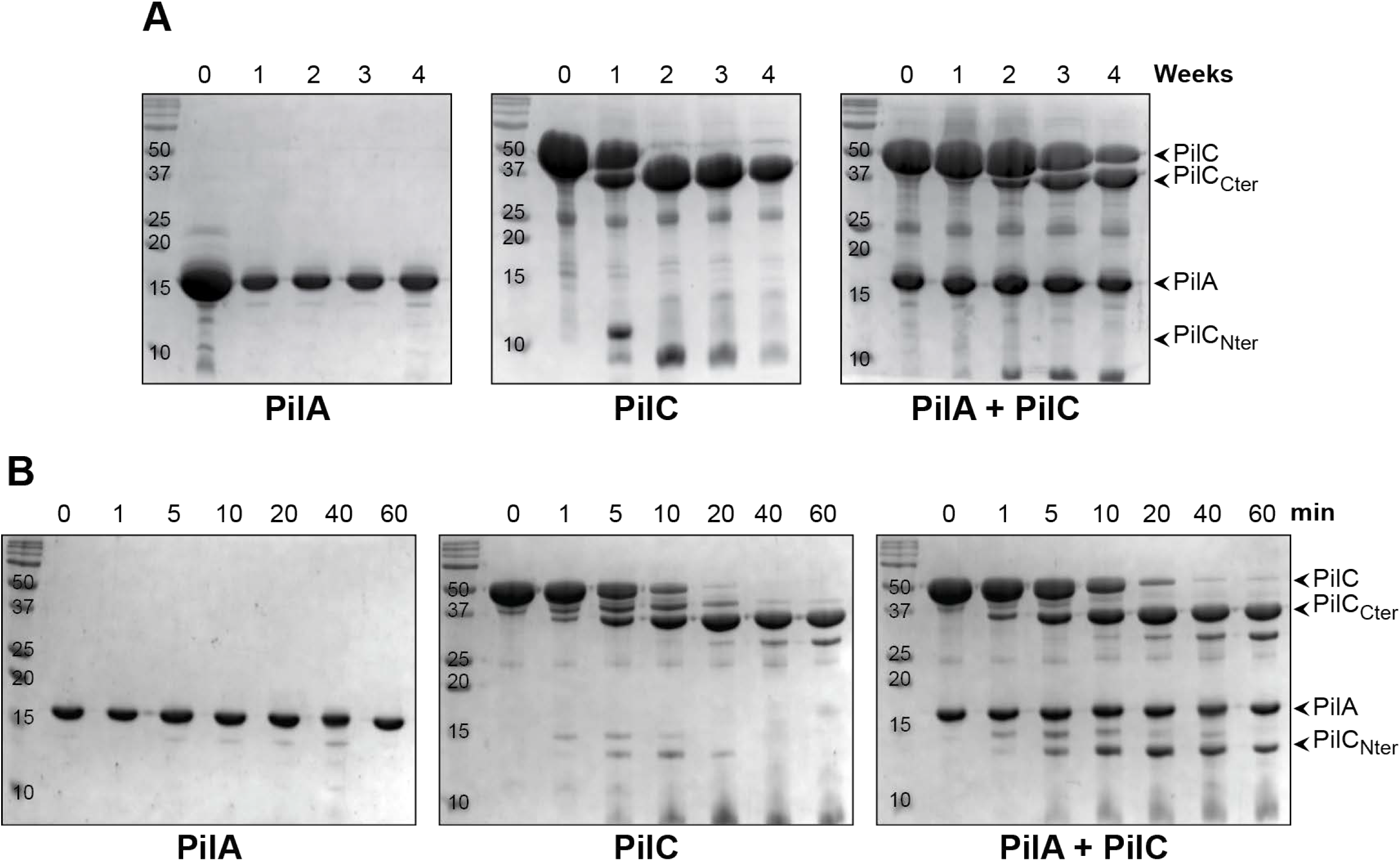
PilA stabilises PilC. **A**) Long-term protein stability experiments. Purified PilA and PilC were mixed in equimolar amounts, and the PilAC complex was isolated by SEC. The complex, as well as PilA and PilC on their own, were kept at 4°C for a month. Samples were taken once a week and analysed by SDS-PAGE/Coomassie. The degradation products (PilC_Nter_ and PilC_Cter_) are identified on the right side. Three independent experiments were performed yielding same results, representative gels are shown. The molecular masses are in kDa. **B**) Trypsin sensitivity assays. Purified PilA, PilC and PilAC were incubated with trypsin (1:1,000 dilution) for 60 min on ice. Samples were taken at various times and analysed by SDS-PAGE/Coomassie. The degradation products (PilC_Nter_ and PilC_Cter_) are indicated on the right side. Three independent experiments were performed yielding similar results, representative gels are shown.

The stabilising effect of PilA on PilC was further characterised by performing trypsin sensitivity assays. We compared trypsin sensitivity of 6His-PilA and 6His-PilC on their own, and in the PilAC complex. In brief, we incubated proteins with trypsin on ice, took samples at 1, 5, 10, 20, 40 and 60 min, which we analysed by SDS-PAGE/Coomassie (Fig. 5B). 6His-PilA was intact over the course of the assay, confirming its high stability. In contrast, 6His-PilC proteolysis began within the first minute and by 20 min the protein was entirely converted into PilC_Nter_ and PilC_Cter_ degradation products. As above, the pilin moiety PilC_Nter_ was completely degraded and no longer detectable after 40 min. In contrast, when it was part of a complex with 6His-PilA, 6His-PilC degradation was delayed since it was still detectable at 20 min (Fig. 5B). In addition, PilC_Nter_ was also stabilised as it was still detectable after 60 min. This indicates that PilA stabilises the pilin moiety of PilC.

Taken together, these findings show that the functional consequence of the PilAC interaction is that PilA stabilises PilC, especially its pilin moiety.

### Crystal structures of PilC reveal a modular pilin

To confirm the prediction that PilC might be a modular pilin, we solved its 3D structure by X-ray crystallography. Due to the instability of its pilin moiety (Fig. 5), we produced a 41.4 kDa recombinant 6His-PilC_Δpilin_ protein in *E. coli*, in which the pilin moiety was missing (Fig. 1C). This protein crystallised readily, and we collected a complete dataset in the space group *P*3_2_21 (Table 1). After phase determination using SeMet crystals, we solved a high-resolution structure at 1.45 Å. Strikingly, while bioinformatics only predicted that the C-terminus of PilC corresponds to a lectin module, our structure reveals two distinct modules (Fig. 6A). The first module adopts an Ig-like fold, consisting of two β-sheets packed against each other (Fig. 6B). This is a widespread domain found in proteins of different functions^39^, which is thought to be involved in protein-protein interactions. Therefore, the Ig-like module in PilC (PilC_Ig_) shows significant structural similarity to multiple structures in the PDB, including the N-acetylglucosamine-binding protein GbpA from *Vibrio cholerae*^40^ (PDB 2XWX). The Ig-like domains in these two proteins, which display no sequence homology, superpose with an RMSD of 2.17 Å (Fig. 6B).

**Fig. 6.**
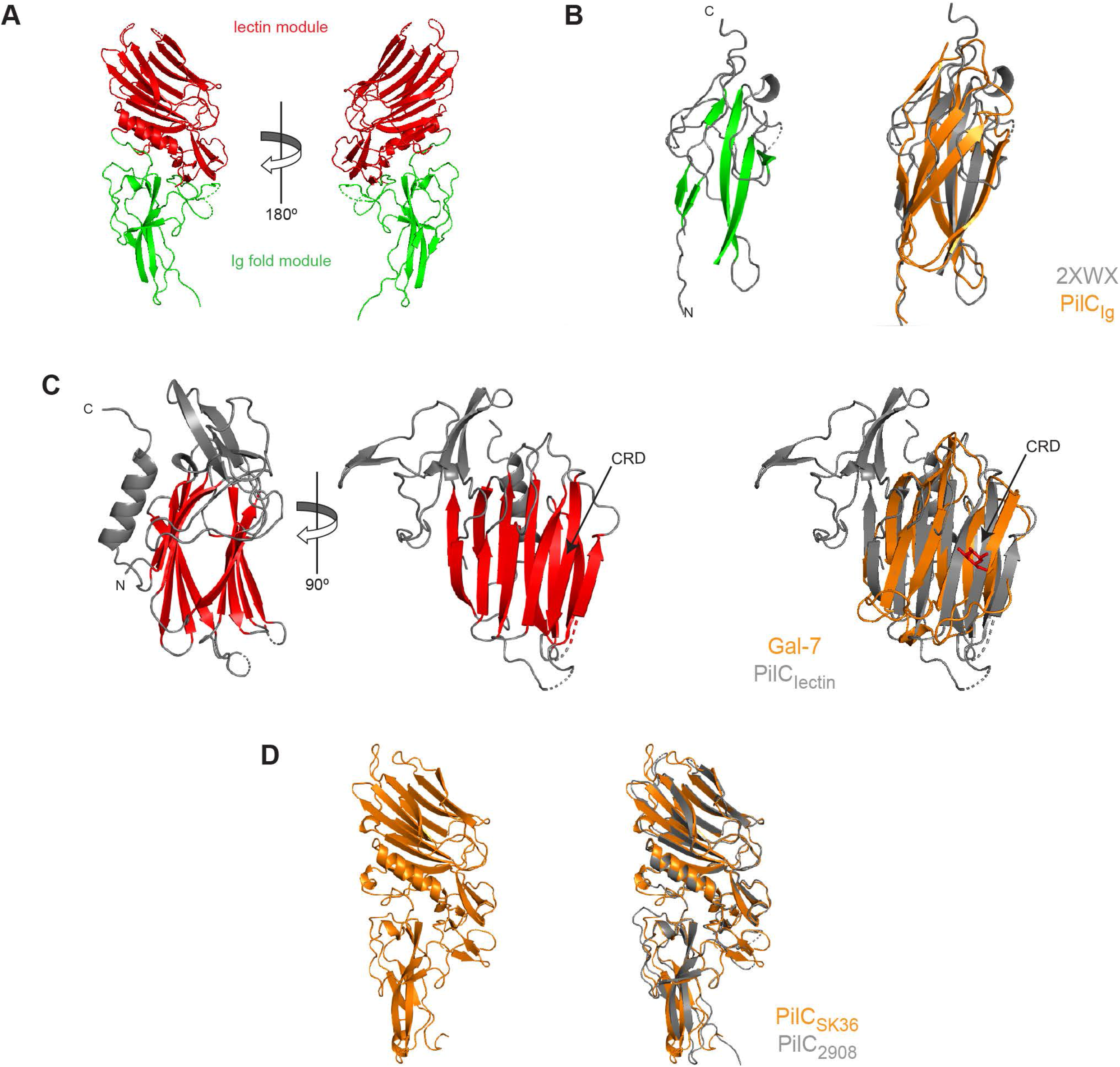
Crystal structures of PilC. **A**) Cartoon views at 180° rotation of the structure of PilC_Δpilin_ from strain 2908 in which two distinct modules have been identified highlighted in green (Ig-like fold) and red (lectin). **B**) Left, Close-up view of the Ig-like fold module (PilC_Ig_). The two β-sheets packed against each other have been highlighted in green. Right, superposition of PilC_Ig_ with the Ig-like fold domain in *V. cholerae* colonisation factor GbpA (PDB 2XWX)^40^. **C**) Left, orthogonal cartoon views of the lectin module (PilC_lectin_) in which the β-sandwich of two opposing antiparallel β-sheets is highlighted in red. The putative CRD is indicated by an arrow. Right, superposition of PilC_lectin_ with human Gal-7 (PDB 2GAL)^41^. The bound galactose in the Gal-7 structure is highlighted in red. **D**) Left, structure of PilC from *S. sanguinis* SK36. Right, superposition of PilC^2908^ and PilC^SK36^. These two proteins, which share 57 % sequence identity (Fig. S4), superpose almost perfectly with an RMSD of only 0.79 Å.

The second module in PilC_Δpilin_, which corresponds to the lectin module (PilC_lectin_), is a β-sandwich characterised by two opposing antiparallel β-sheets, each composed of six strands (Fig. 6C). The β-sheets yield one concave and one convex side. Confirming the bioinformatic predictions, PilC_lectin_ shows significant structural similarity to multiple carbohydrate-binding proteins, including galectins such as human Gal-7 (PDB 2GAL)^41^. PilC_lectin_ and Gal-7, which display no sequence homology, superpose with an RMSD of 1.95 Å (Fig. 6C). This similarity is interesting because the ligand-binding ability of galectins is well-characterised. In the Gal-7 structure in complex with galactose (Fig. 6C), the carbohydrate-recognition domain (CRD) forms a patch on the concave side of the protein^41^. Since there is good structural homology in that part of PilC_lectin_ (Fig. 6C), we speculate that the CRD in PilC might be on the concave side of PilC_lectin_.

To try to determine a PilC structure encompassing the pilin moiety as well, we expressed and crystallised the corresponding protein from another *S. sanguinis* strain SK36 (PilC^SK36^), in which the N-terminal 33 residues were replaced by 6His. We collected a complete dataset in the space group *P*2_1_2_1_2_1_ (Table 1) and solved a 1.60 Å resolution structure. Surprisingly, although the purified protein contained the pilin moiety, this portion was missing in the crystals. The pilin moiety in PilC^SK36^ therefore became degraded during prolonged incubation in the crystallisation trays, which is likely to be due to an inherent instability. When we compared our two PilC structures, we found that they are essentially identical, superposing onto each other with an RMSD of only 0.79 Å (Fig. 6D). This suggests that these proteins – showing only 57 % sequence identity (Fig. S4) – are likely to bind similar glycan ligands.

### PilC binds two types of glycans prevalent in the human glycome

The presence of a lectin domain suggests that PilC might be an adhesin recognising host glycans. We therefore investigated the glycan binding ability and specificity of PilC using a glycan microarray with oligosaccharide probes immobilised using the neoglycolipid (NGL) technology^42^. Microarray analysis with 6His-PilC revealed that it binds to a range of glycan probes – results are shown for 672 sequence-defined oligosaccharide probes (Supplementary dataset 1) – corroborating its lectin activity (Fig. 7A). Two major types of negatively charged glycans were bound: sialylated glycans and sulphated glycosaminoglycans (GAG). Most of the sialylated glycan ligands display a α2-3-sialyl linkage and terminate with 3′-sialyllactose (3’-SL, NeuAc(α2-3)Gal(β1-4)Glc) or 3′-sialyl-N-acetyllactosamine (3’-SLN, NeuAc(α2-3)Gal(β1-4)GlcNAc), which are similar trisaccharides. Also well recognised (Fig. 7A) were ganglioside GD1a and four probes terminating with the Sd^a^ antigen NeuAc(α2-3)[GalNAc(β1-4)]Gal(β1-4)GlcNAc(β1-3)Gal, which is known to be present in human secretions including saliva^43,44^. A few probes displaying other sialyl linkages – including α2-9-polysialic acid – were also bound (Fig. 7A). The sulphated GAG ligands are mainly oligosaccharide probes derived from heparin, representative of the highly sulphated domains of heparan sulphate, which is ubiquitously expressed on the cell surface and in the extracellular matrix. Both sialylated glycans and GAGs are prevalent in the human glycome^45^.

**Fig. 7.**
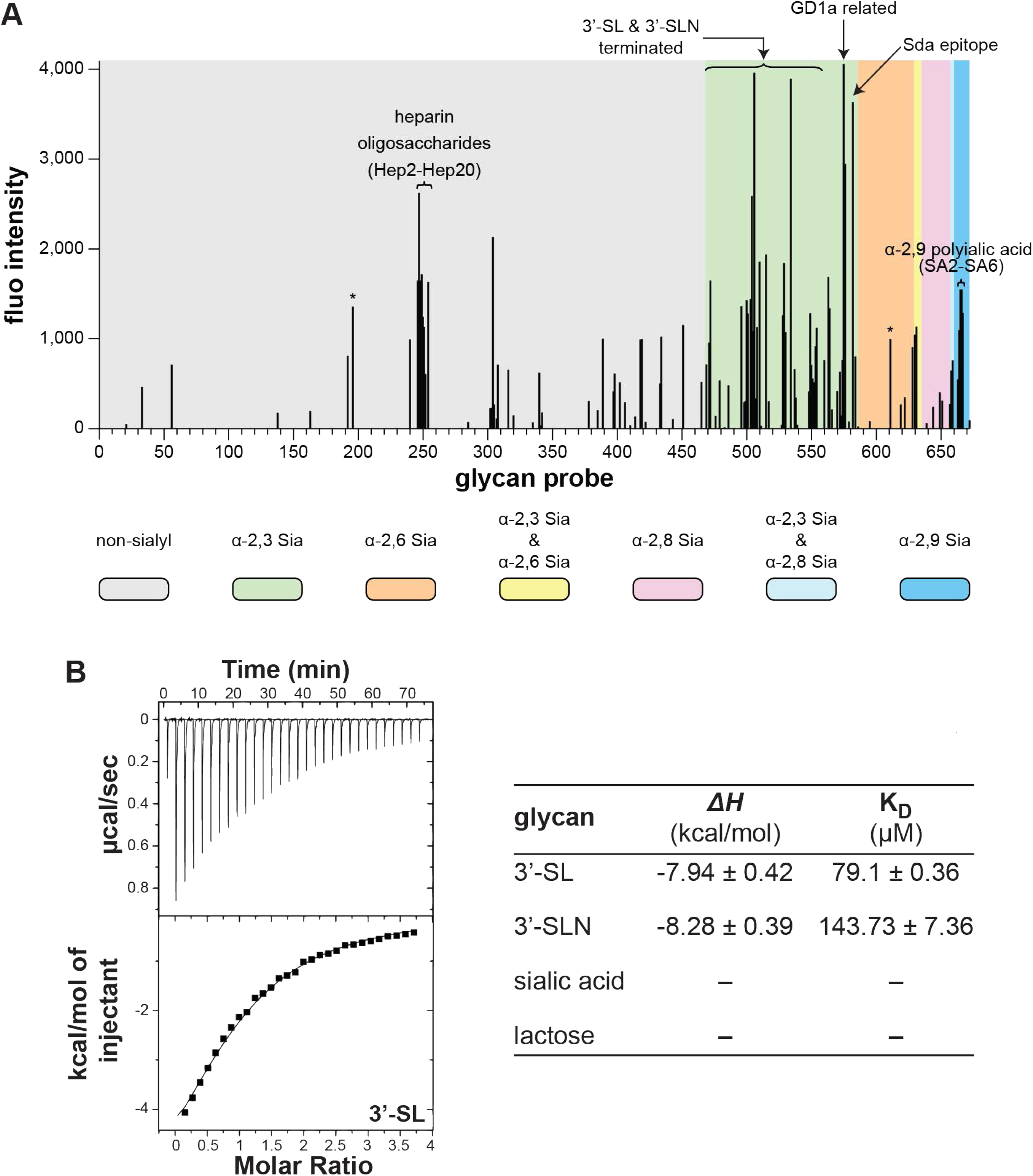
PilC specifically binds sialylated glycans and GAG. **A**) Glycan microarray screening analyses of 6His-PilC, as a protein-antibody pre-complex. Multiple experiments were performed yielding similar results. Results of one representative experiment are shown as the means of fluorescence intensities of duplicate spots of glycan probes, printed at 5 fmol/spot. The 672 lipid-linked probes are grouped according to sialyl linkages as annotated by the coloured panels. The full list of glycan probes, their sequences, and binding scores are given in Supplementary dataset 1. * Signals with large error bars due to artefacts on the array slides. **B**) ITC quantification of PilC_Δpilin_ affinity for sialylated glycans. The calculated K_D_ are the average ± SD from three independent experiments. No binding was seen for sialic acid and lactose. For binding to 3’-SL, representative ITC titration curve (upper panel) and binding isotherm (lower panel) are presented.

We tested whether the lectin activity of PilC is affected by its pilin moiety, by probing the glycan microarray with 6His-PilC_Δpilin_, the purified protein without pilin moiety. We found that PilC_Δpilin_ bound the same type of glycans as PilC, suggesting that glycan binding is largely due to PilC’s lectin module. However, the pilin moiety affected the glycan binding characteristics (Fig. S5). When compared to PilC, PilC_Δpilin_ binds strongly to heparin oligosaccharide probes and weakly to a more restricted range of α2-3-linked sialyl glycans (Fig. S5). In parallel, we observed for PilC^SK36^ a similar binding profile as for PilC, showing that the glycan-binding ability is conserved in PilC orthologs (Fig. S6). Next, using ITC, we quantified the affinity of PilC_Δpilin_ for 3’-SL, 3’-SLN and their constituent subunits (sialic acid and lactose) (Fig. 7B). No binding was seen with lactose or sialic acid. In contrast, PilC bound both 3’-SL and 3’-SLN, in accord with the microarray findings. There was a slight preference for 3’-SL over 3’-SLN, with K_D_ of 79.1 ± 0.36 µM and 143.73 ± 7.36 µM, respectively (Fig. 7B).

Finally, we probed the glycan-binding activity of PilC by mutagenesis of residues on the concave side of PilC_lectin_ where we the CRD is predicted to be (Fig. 6C). We made series of PilC_Δpilin_ derivatives in which surface-exposed residues in the putative CRD were changed by site-directed mutagenesis (Fig. 8A). We then purified and quantified the affinity of these mutant proteins for 3’-SL using ITC and compared it to WT

**Fig. 8.**
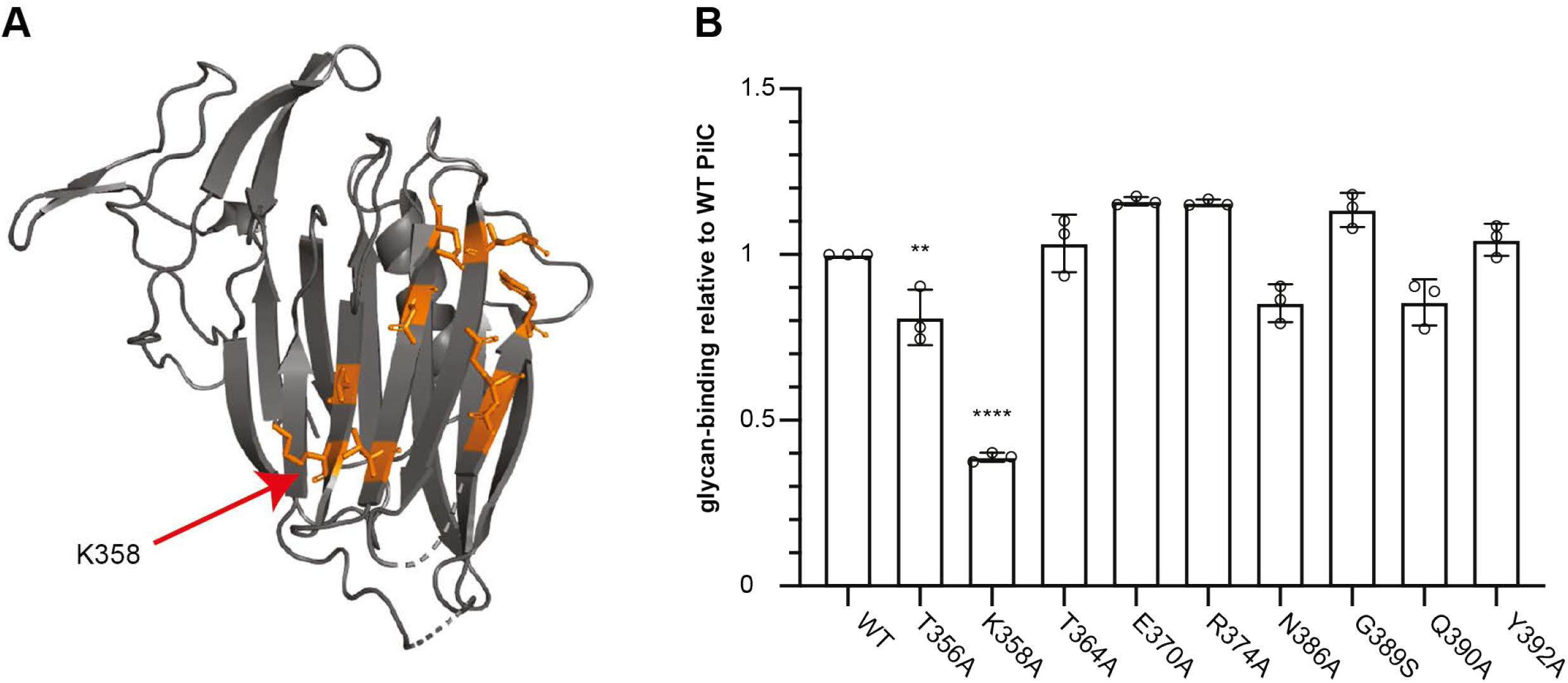
Probing the putative carbohydrate-recognition domain in PilC by mutagenesis. **A**) Cartoon view of PilC_lectin_ where the residues in the putative CRD – targeted by SDM – are highlighted in orange. **B**) ITC quantification of binding to 3’-SL by PilC_Δpilin_ mutant proteins. The results, presented as K_D_ ratios to the WT, are the average ± SD from three independent experiments. Statistical significance was calculated using one-way ANOVA followed by Dunnett’s multiple comparison tests.

PilC (Fig. 8B). This revealed that two of the mutants PilC_T356A_ and PilC_K358A_ displayed significantly reduced affinity for 3’-SL, which was particularly pronounced for PilC_K358A_, with 39.3 ± 1.4 % of the WT binding. This finding indicates that a portion of the concave side of its lectin module is indeed involved in the ability of PilC to bind specific glycans.

### Modelling predicts that the PilAC complex is located at the tip of *S. sanguinis* T4P, together with PilB

Since the instability of PilC pilin moiety (Fig. 5) precluded us from determining a crystal structure of the PilAC complex, we tried to model this complex using AlphaFold^38^. We first modelled PilC_pilin_, generating a prediction of high quality (Fig. S7). The globular head of PilC is predicted to display a highly unusual β-sheet composed of three anti-parallel β-strands, where the orthogonal β1 and β3 are linked by a kinked β2 (Fig. 9A). We then modelled the PilAC_pilin_ complex (Fig. 9B). PilA is predicted to be on the top, interacting with PilC_pilin_ with an axial rise that might correspond to the axial rise in *S. sanguinis* T4P. PilA extends the β-sheet of PilC, which is likely to explain the stabilising effect of PilA on PilC (Fig. 9B). The complex is predicted to be stabilised by a series of 21 hydrogen bonds and four salt bridges (Fig. S8). The predicted PilAC interaction interface (Fig. S8) is in strikingly good agreement with that identified by NMR (Fig. 4B), including the α1 and α2 helices, and the β1 and β2 strands of PilA (Fig. S8). This significantly strengthens the validity of the model.

**Fig. 9.**
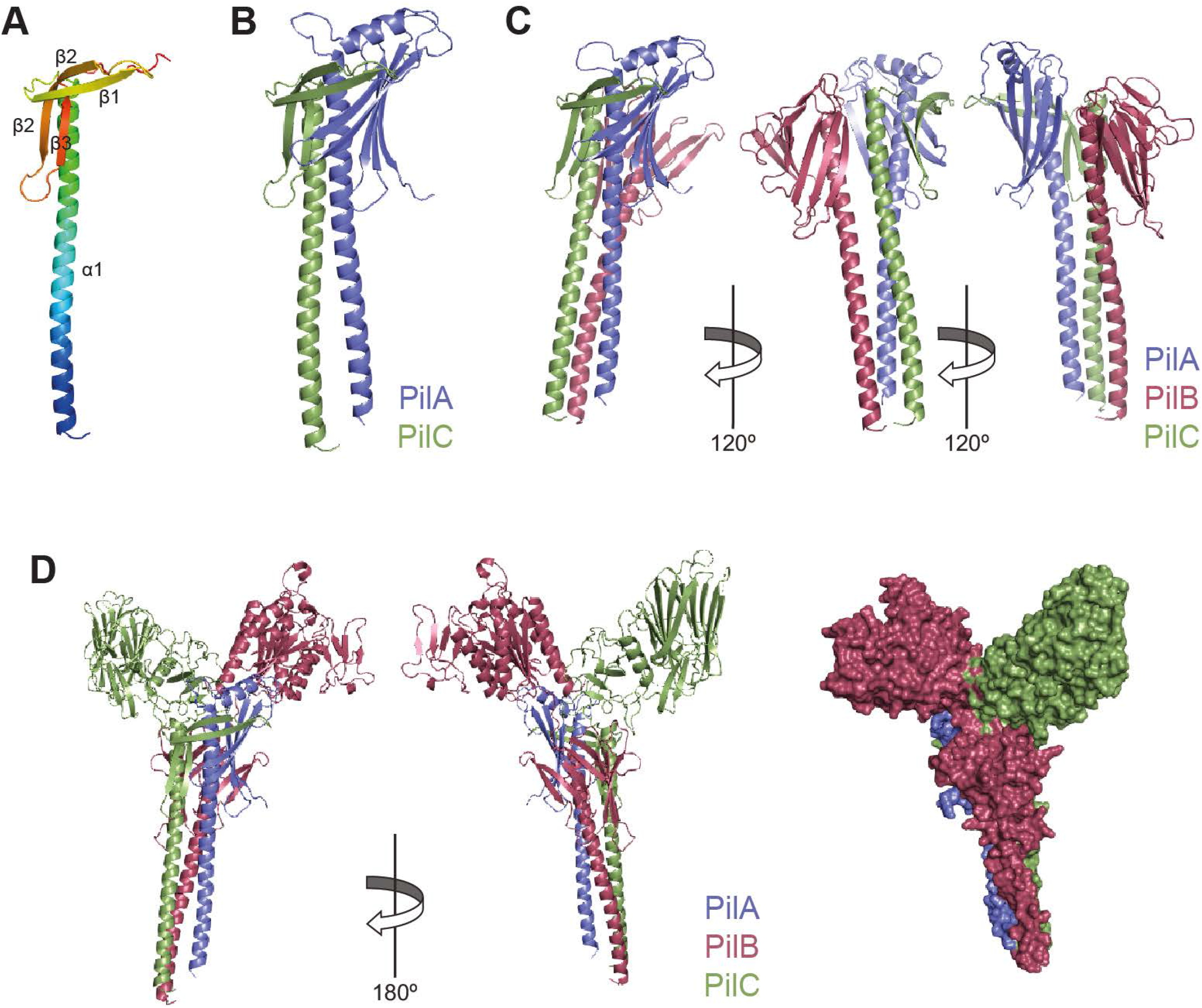
Modelling the complex of minor pilins in *S. sanguinis* T4P. **A**) Cartoon view of the AlphaFold model of PilC_pilin_ rainbow-coloured from blue (N-terminus) to red (C-terminus). This highlights an unusual globular head with a three-stranded β-sheet, where a heavily kinked β2 links β1 and β3, which are orthogonal. **B**) Model of the PilAC_pilin_ complex. PilA interacts with PilC_pilin_ by extending its β-sheet, which is likely to explain the stabilising effect of PilA on PilC. The axial rise between PilA (on top) and PilC might correspond to the axial rise in *S. sanguinis* T4P. **C**) Model of the PilAC_pilin_B_pilin_ complex. Cartoon views at 120° rotation show a quasihelical complex with the same axial rise as in PilA-PilC_pilin_. **D**) Model of the PilACB complex. Left, cartoon views at 180° rotation. Right, surface representation. Although PilA is added first, the extra modules in PilB and PilC are capping the complex like “open wings”. For this reason, the PilACB complex could only be accommodated at the tip of the T4P.

The other modular pilin in *S. sanguinis* T4P, PilB, was previously characterised and predicted to be at the pilus tip^6^. We therefore wondered if a co-existence of PilB with the PilAC complex was possible, which we assessed by modelling using AlphaFold. As above, we first modelled a putative PilAC_pilin_B_pilin_ complex (Fig. 9C), which predicts that a quasihelical complex is possible with the same axial rise seen in PilAC_pilin_. In this complex, PilA is predicted to be on the top, interacting with PilC, which itself interacts with PilB (Fig. 9C). We then modelled the full-length PilACB complex using AlphaFold. Since the prediction for full-length PilC is of poor quality, especially for the Ig fold module, we produced a full-length PilC model by pasting the predicted PilC_pilin_ onto our crystal structure where the Ig fold module is well resolved. The model suggests that the two modular pilins PilB and PilC can be accommodated in the same complex despite their bulky extra domains (Fig. 9D). Some level of movement for the extra modules in PilB and PilC is expected because the short loops that connect them to their respective pilin moieties are likely to be flexible. Although PilA is predicted to be added first, the extra modules in PilB and PilC are capping the complex and look like “open wings”. Due to this peculiar architecture, the PilACB complex could only be accommodated at the tip of the T4P, as previously suggested for HIJK^25^. This is likely to optimise the presentation of the adhesive modules in PilB and PilC, promoting *S. sanguinis* adhesion to host surfaces.

## Discussion

T4F are a superfamily of filamentous nanomachines ubiquitous in bacteria and archaea^1,2^, mediating a wide variety of functions ranging from adhesion, to motility, via protein secretion. Although T4F have been studied for 40 years – notably their T4P archetype in human diderm bacterial pathogens^46^ – many aspects of their complex biology remain poorly understood. This holds especially true for the filament – the “business end” of T4F nanomachines – where the roles of all pilin subunits are yet to be understood. Therefore, in this report we used the inherent simplicity of T4P in monoderm bacteria – which recently opened new research avenues^30^ – to complete a structure/function analysis of all the pilins in *S. sanguinis*^31^ T4P. This led to the notable findings discussed below, shining new light on T4F.

The first important finding in this study – with widespread implications in T4F biology – is the unexpected discovery that the minor pilin PilA from *S. sanguinis* is a structural homolog of the I subunit of the widely conserved HIJK minor pilins. The broader than predicted conservation of that minor pilin is a testament to the central role it plays in T4F biology. There are multiple analogies between the two systems, suggesting that PilABC is a rudimentary, perhaps ancestral, version of the HIJK complex. The HIJK pilins are found in different T4F where they play conserved role since they are functionally interchangeable^24^. These pilins are encoded by an operon, and the K subunit lacks the usually conserved Glu residue in fifth position of the mature pilin^4^. PilABC pilins are also encoded by an operon, and PilA lacks the Glu_5_ residue^35^. Structural studies showed that HIJK pilins interact to form a quasihelical complex^25,26^ in the order KIJH, from top to bottom. The complex – capped by the modular K subunit – was thus proposed to be at the tip of T4F because no additional subunits could fit above the bulky K without significant steric hindrance^25^, which was recently confirmed by cryo-ET^27^. Similarly, we found that PilA and PilC interact via their pilin moieties, forming a complex likely to be at the tip of *S. sanguinis* T4P because no additional subunits could fit above the bulky PilC modular pilin. Since PilB is another bulky modular pilin predicted to be tip-located^6^, we propose that PilA, PilB, PilC pilins interact to form a quasihelical complex in the order PilACB, from top to bottom. Although PilA is added first, the extra modules in PilB and PilC cap the PilABC complex like open wings. Due to this peculiar architecture – resembling the Winged Victory of Samothrace, a masterpiece of Greek sculpture – the PilABC complex could only be accommodated at the tip of the T4P. Critically, both HIJK^21–23^ and PilABC^34^ pilins are essential for filament assembly, which is compatible with PilA being a nucleator initiating filament assembly, a role attributed to the I subunit in the HIJK complex^28,29^. For the HIJK complex, it was proposed that (1) I interacts with J in the membrane to form a staggered complex, (2) the IJ complex interacts with K via its I subunit, initiating filament assembly by forming a quasihelical complex^25^, (3) the IJK complex interacts with H via its J subunit^26^, and (4) the HIJK complex interacts with the major pilin via its H subunit^26^. The series of events is similar for PilABC, albeit simpler. (1) PilA interacts with PilC and stabilises it (it remains to be seen if the I subunit in HIJK plays a similar role), (2) PilC interacts with PilB, and (3) the PilABC complex most likely interacts with the major pilins via its PilB subunit. In both systems, the I/PilA subunit initiates filament assembly and facilitate the presentation of “effectors” at the tip of filaments. In T4P, the effectors are either the two modular pilins PilB and PilC with adhesive properties (in *S. sanguinis*), or the non-pilin adhesin PilC/PilY1 (in diderm model species). Interestingly, cryo-ET analysis of *M. xanthus* T4P revealed that PilC/PilY1 forms a kinked tip structure^27^, which looks like one of the open wings in PilABC. It is possible to speculate that in T2SS, the HIJK complex might be involved in recognising secreted substrates^47^.

Drawing a picture of the roles of all the pilin subunits in a model T4F system is the second major achievement in this study. *S. sanguinis* T4P are composed of five pilins: two major (PilE1, PilE2) and three minor (PilA, PilB, PilC) ones. PilE1 and PilE2 – 85 % identical in sequence^34^ – are the two major pilins. PilE1 and PilE2 follow universally shared principles for processing by the PPase and display lollipop structures^35^. This suggests that *S. sanguinis* T4P are canonical T4F, *i.e.*, helical polymers where pilins pack within the core of the filament and are held together by extensive interactions between their α1-helices^11^. Although unusual – T4F are most often homopolymers of only one major pilin – the heteropolymeric composition of *S. sanguinis* T4P with two major pilins in nearly equal amounts^35^ is not unique. Recently, the structure of a heteropolymeric archaeal T4F used for swimming has been elucidated by cryo-electron microscopy^16^ (cryo-EM). The archaellum from *Methanocaldococcus villosus* is composed of two alternating major pilin subunits^16^ (ArlB1, ArlB2). It was proposed that the filament is assembled by pre-formed ArlB1-ArlB2 heterodimers, which may be the fundamental building block of the *M. villosus* archaellum^16^. Whether a similar alternating arrangement of PilE1 and PilE2 exists within the *S. sanguinis* T4P, could be tested by cryo-EM. PilA, PilB and PilC are the three minor pilins in *S. sanguinis* T4P, which can be divided into two categories: non-modular (PilA) and modular (PilB, PilC) pilins. We have now defined their respective roles. PilA, which displays a canonical lollipop structure, interacts strongly and exclusively with PilC. Pull-down experiments suggest that PilA and PilC interact via their pilin moieties, which was confirmed by NMR. The interaction interfaces determined by NMR and predicted by modelling are in strikingly good agreement. As a functional consequence, PilA dramatically stabilises the PilC pilin moiety, which is highly unstable on its own. Modelling suggests that this is done by β-strand complementation. PilA thus acts as an intra-pilus chaperone, protecting PilC from proteolysis by perhaps participating in its proper folding. In addition, it is likely that PilA also participates in the proper localisation of PilC at the pilus tip. We previously showed that PilB displays a modular architecture with a bulky functional module grafted onto a pilin moiety^6^. The vWA (IPR036465) module in PilB makes it a *bona fide* adhesin, playing a key role in *S. sanguinis* adhesion to host cells by binding protein ligands such as fibronectin and fibrinogen^6^. Since *S. sanguinis* T4P are critical for disease progression^48^, these findings are consistent with PilB-mediated adhesion being involved in IE. Here, we show that PilC is also a modular pilin with two extra modules, an Ig-like fold (not revealed by bioinformatics) and a ConA-like lectin/glucanase module (SSF49899). While the function of the Ig-like module remains unclear, we showed that the ConA-like module in PilC is a *bona fide* lectin. PilC binds specifically to two major types of glycans – (1) sialylated glycans terminated with α2-3-sialyl linkage as in 3’-SL or 3’-SNL, ganglioside GD1a or glycans carrying the Sd^a^ epitope, and (2) heparin oligosaccharide probes representative of sulphated GAGs – both prevalent in the human glycome^45^. The measured affinities for 3’-SL and 3’-SLN – in the µM range – are consistent with those previously determined for other SSF49899-containing proteins^49^. Affinities could be much higher for larger glycan determinants, or when there is multivalent presentation (abundance) of a glycan ligand at the cell surface^45,49^. The physiological ligands for PilC are yet to be defined – identifying ligands for lectins is often difficult^45,50^ – but it is likely that this modular pilin is involved in *S. sanguinis* commensalism or IE. Indeed, 3’-SL or 3’-SLN terminated glycans are abundant in the oral cavity^32^, especially on heavily glycosylated salivary mucins^45^. Interestingly, α2-3-sialyl glycans have been reported to be ligands of the Siglec-like domain of serine-rich repeat adhesins expressed by oral streptococci^51^. The full picture of *S. sanguinis* T4P that has now emerged is the following. *S. sanguinis* T4P are heteropolymeric filaments composed of PilE1 and PilE2, with helical symmetry parameters that are yet to be determined, capped by a complex of three minor pilins (PilA, PilB, PilC) mediating bacterial adhesion to different eukaryotic receptors. The adhesive modules in PilB and PilC are ideally placed to maximise adhesion. However, although this is less likely, we cannot exclude the possibility that *S. sanguinis* expresses two distinct pili, capped either by PilB or PilAC.

In conclusion, by providing perhaps the most detailed structure/function characterisation of the role of all pilins in a model T4P, this study sheds light on important aspects of T4F biology. The finding that PilABC might be a rudimentary version of the widely conserved HIJK complex strengthens the notion that complexes of minor pilins capping T4F facilitate the presentation of different effectors at the tip of filaments, thus directly promoting T4F exceptional functional versatility^1^. Our findings have general implications for T4F and pave the way for further investigations that will improve our understanding of these fascinating filaments.

## Material and methods

### Strains and growth conditions

*E. coli* strains were grown in liquid or solid lysogeny broth (LB) medium (Difco), containing 50 µg/ml kanamycin (Sigma) when required. The strains and plasmids used in this study are listed in Table S1. *E. coli* DH5α was used for cloning, while *E. coli* BL21(DE3) was used for routine protein expression and purification. For purification of proteins labelled with SeMet, *E. coli* B834(DE3) was grown in chemically defined medium (CDM) supplemented with 20 mg/ml SeMet (Sigma).

Chemically competent cells were prepared as described elsewhere^52^. DNA manipulations were performed using standard molecular biology techniques^53^. All PCR were done using high-fidelity DNA polymerases (Agilent). The primers used in this study are listed in Table S2. Missense mutations in recombinant proteins were generated by QuikChange site-directed mutagenesis (Agilent) using the respective pET28b derivatives as templates. To purify PilA and PilC, the soluble portion of these proteins (excluding the leader peptide and the predicted α1N helix) was fused to non-cleavable N-terminal 6His or Strep II tags and cloned into pET-28b (Novagen). We amplified *pilA* directly from the *S. sanguinis* 2908 genome, while *pilB* and *pilC* were synthetic genes codon-optimised for expression in *E. coli* (GeneArt).

### Protein purification

The recombinant proteins were purified in two steps: affinity chromatography and SEC. The respective pET-28b derivatives were transformed into *E. coli* BL21(DE3). A single colony was picked and grown O/N in LB with kanamycin. The following morning, this culture was back diluted 1/100 in 1 l of the same medium. Once the OD_600_ (optical density at 600 nm) reached 0.8-1, the cultures were cooled down to 16°C, and protein expression was induced O/N with 0.5 mM isopropyl 1-thio ß-D-galactopyranoside (IPTG) (Merck Chemicals). The next day, the cells were harvested by centrifugation at 6,000 *g* for 20 min, and the pellets were frozen at −80°C in the appropriate binding buffer, containing the SigmaFast EDTA-free protease inhibitor cocktail (Sigma). Cells were lysed by repeated cycles of sonication (5 sec on/off pulses for 3-5 min) and the clarified cell lysate was prepared by centrifugation at 11,000 *g* for 30 min.

6His-tagged proteins were affinity-purified using Gravity Flow Chromatography columns (Bio-Rad) loaded with 1-2 ml Ni-NTA agarose resin (Qiagen) pre-equilibrated with binding buffer. The resin was mixed with the cell lysate and the column was allowed to drain. The column was then washed with binding buffer (50 mM HEPES pH 7.5, 150 mM NaCl, 20 mM imidazole) several times, before being eluted with elution buffer (50 mM HEPES pH 7.5, 150 mM NaCl, 300 mM imidazole). Strep-tagged proteins were affinity-purified on an ÄKTA Purifier using 5 ml StrepTrap HP columns (GE Healthcare) according to manufacturer’s instructions. The columns were washed with binding buffer (50 mM HEPES pH 7.5, 150 mM NaCl) and protein was eluted with elution buffer (50 mM HEPES pH 7.5, 150 mM NaCl, 2.5 mM desthiobiotin). Both the 6His- and Strep-tagged proteins were then further purified by SEC on an ÄKTA Purifier using a Superdex 200 16/600 GL column (GE Healthcare), and simultaneously buffer-exchanged into 50 mM HEPES pH 7.5, 150 mM NaCl. Protein concentration was quantified spectrophotometrically on a NanoDrop Lite (Thermo Fisher Scientific).

To purify SeMet-labelled PilA and PilC for phasing, the corresponding pET-28b derivatives were transformed in *E. coli* B834(DE3). A single colony was picked and grown O/N in LB with kanamycin. After back dilution, transformants were grown at 37°C in selective liquid LB, until OD_600_ reached 0.6-0.7. The cells were pelleted at 8,000 *g* for 5 min and washed twice with 2 ml of CDM containing no Met. The pellets were then washed with 2 ml of CDM supplemented with 20 mg/ml L-Met (Sigma) and used to inoculate, at 1/200 dilution, 20 ml of CDM supplemented with 20 mg/ml Met. These cultures were grown O/N at 37°C. Cells were pelleted and washed three times with CDM. Then, the pellets were re-suspended in 20 ml of CDM, supplemented with 20 mg/ml SeMet, and used to inoculate 1 l of CDM supplemented with SeMet. Cells were grown at 37°C until OD_600_ reached 0.5-0.7. The cultures were cooled down to 16°C, and protein expression was induced by adding 1 mM IPTG and 4 ml of 36 % glucose (w/v). The cultures were further supplemented with another 4 ml of 36 % glucose 2.5 h later. The next day, the cells were harvested, and SeMet-labelled proteins were purified as above.

To purify ^15^N and/or ^13^C-labelled PilA for NMR analysis, the corresponding pET-28b derivative was transformed in *E. coli* BL21(DE3). A single colony was used to inoculate a starter culture, which was back-diluted 1/100 into 1 l kanamycin-supplemented LB the following morning. The cultures were grown at 37°C until the OD_600_ reached 0.9. The cells were harvested by centrifugation at 6,000 *g* for 10 min at 4°C. The pelleted cells were resuspended in 490 ml M9 salts (3.37 mM Na_2_HPO_4_. 2.2 mM KH_2_PO_4_, 0.855 mM NaCl, pH 7.2), containing 1 ml 1 M MgSO_4_ and 250 µl 0.2 M CaCl_2_. The cell suspension was supplemented with 5 mg vitamin B1, 50 µg/ml kanamycin, 0.5 g ^15^NH_4_Cl and 2 g unlabelled D-glucose (or ^13^C-labelled glucose if double isotopic labelling was required) dissolved in 10 ml H_2_O and filter-sterilised. The cells were rested at 16°C for 20 min before protein expression was induced O/N with 0.5 mM IPTG. The cells were harvested, lysed, and purified as described above. SEC was performed in NMR buffer (10 mM Na_2_HPO_4_/NaH_2_PO_4_ pH 7.0, 150 mM NaCl).

### Protein crystallisation and structure determination

Protein crystals for SeMet-labelled PilA were produced using 0.4 µl sitting drops with a 1:1 ratio protein to mother liquor (0.1 M Tris pH 8, 23 % PEG 2K (w/v), 0.3 M Mg(NO_3_)_2_·6H_2_O). Data was collected at the Diamond beamline i04-1, processed using Xia2^54^, and phased using autoSHARP^55^. Initial structure was produced using CRANK2^56^ and autobuild, with some manual building in Coot^57^. Refinement was performed using Coot and phenix.refine^58^. Structure validation was performed using MolProbity^59^.

Protein crystals for SeMet-labelled PilC were produced using 0.2 µl sitting drops with of 1:1 ratio protein to mother liquor (0.1 M Pipes pH 7 15 % w/v PEG broad smear, 0.2 M (NH_4_)_2_SO_4_, 0.0 1M cadmium chloride hemi (pentahydrate)). Data was collected at the Diamond beamline i03. Data was processed using Xia2 and phased using autoSHARP. Structure was produced using autobuild and refined with manual building in Coot and phenix.refine. MolProbity was used for structure validation.

Protein crystals for PilC^SK36^ were produced in 0.2 µl sitting drop with a 1:1 protein to mother liquor (0.2 M MgCl_2_·6H_2_O, 20 % w/v PEG 3350). Data was collected at the Diamond beamline i04 and processed using Xia2. Structure was produced using Coot and phenix.refine – using the PilC structure from strain 2908 for molecular replacement – and validated using MolProbity.

All the 3D structures determined during this study have been deposited in the PDB and are available under accession codes 7O5Y (PilA), 7OA7 (PilC), and 7OA8 (PilC^SK36^).

### Pull-down assays

Pull-down assays were carried out using Dynabeads His-tag Isolation and Pulldown (Invitrogen). The 6His-tagged soluble pilin domains were used as bait, while Strep-tagged soluble pilins were used as prey. For each pull-down reaction, 25 µl of magnetic beads and 1.5 nmol of protein were used (23 µg for PilA, 75 µg for PilC, and 60 µg PilC_Δpilin_). The pull-down assays were performed three times for each combination. Prior to the pull-down assays, the bait and prey proteins were mixed in 1 ml of binding buffer (50 mM HEPES pH 7.6, 150 mM NaCl, 10 mM imidazole, 0.1 % Tween-20) and incubated on ice for 1 h. Meanwhile, the magnetic beads were incubated with 700 µl blocking buffer (50 mM HEPES pH 7.6, 150 mM NaCl, 10 mM imidazole, 0.1 % Tween-20, 5 % skim milk) on a rotating wheel at 4°C. After 1 h, the blocking reaction was placed on a magnet for 10 sec to capture the beads, and the flow-through was discarded. The beads were then rinsed twice with 700 µl binding buffer and mixed with 1 ml of bait-prey reaction mixture. Following a 20-min incubation on a rotating wheel at 4°C, the beads were captured on the side of the tube using a magnet, and the flow-through was removed. Beads were washed 10 times by vortexing them with 500 µl washing buffer (50 mM HEPES pH 7.6, 150 mM NaCl, 20 mM imidazole, 0.1 % Tween-20) for 10 sec. After the final wash, the beads were incubated with 100 µl elution buffer (50 mM HEPES pH 7.6, 300 mM NaCl, 500 mM imidazole, 0.1 % Tween-20) for 5 min on a rotating wheel at 4°C. The beads were captured on the side of the tube using a magnet, and the flow-through was carefully transferred to a fresh tube. For each pull-down reaction, input, flow-through, and elution samples were analysed. The samples were mixed with 2x Laemmli Buffer (Bio-Rad), boiled for 5 min at 100°C, and subsequently analysed by immunoblotting.

### SDS-PAGE and immunoblotting

SDS-PAGE was carried out using 1x Tris/Glycine/SDS Buffer (Bio-Rad) in a Mini-Protean Tetra cell system (Bio-Rad) for 1 h at 200 V. The Precision Plus Protein All Blue Prestained Protein Standards (Bio-Rad) was used as molecular weight marker and loaded alongside the protein samples. Gels were either stained with Bio-Safe Coomassie (Bio-Rad) and imaged with a Gel Doc EZ Imager (Bio-Rad) or transferred to a membrane and analysed by immunoblotting.

Immunoblotting was done as follows. After proteins were separated by SDS-PAGE, they were transferred onto Amersham Hybond ECL nitrocellulose membrane (GE Healthcare). The wet transfer was carried out for 1 h at 100 V in ice-cold buffer (39 mM glycine, 48 mM Tris base, 0.037 % SDS, 20 % isopropanol). The blotted membranes were blocked for 1 h at room temperature, while shaking, in PBS supplemented with 0.1 % Tween-20 (PBST) containing 5 % (w/v) skim milk powder (VWR). The membranes were then incubated for 1 h with primary antibodies diluted 1/3,000 in 5 % milk PBST. The specific antibodies generated in rabbits against PilE1, PilE2, PilA, PilB, PilC, were previously described^34,35^. Following three 10-min washes with PBST, the membranes were incubated for 1 h with an anti-rabbit secondary antibody conjugated to horseradish peroxidase (GE Healthcare), diluted 1/10,000 in PBST. The membranes were washed again three times for 10 min in PBST, dried and developed with Amersham ECL Prime Western Blotting Detection Reagent (GE Healthcare). Protein bands were detected using a ChemiDoc Imaging System (Bio-Rad).

### SEC-MALS

SEC-MALS was performed using an ÄKTA Prime system with a S200 10/300 GL column (GE Healthcare) and a MALS detector (Wyatt). The column and the system were equilibrated in buffer (50 mM HEPES pH 7.5, 150 mM NaCl) for 48-72 h to minimise and stabilise the light scattering and background noise. Protein samples of 150 µl (45 µM PilA, 50 µM PilC, 40 µM PilA-PilC) were loaded onto the column at 0.2 ml/min. The UV, light scattering and refractive index signals were analysed with the ASTRA chromatography software.

### ITC

ITC was used to study the interaction between the minor pilins PilA and PilC, as well as the binding of PilC_Δpilin_ to sugar ligands 3’-SL and 3’-SLN. The experiments were performed on a MicroCal ITC200 Malvern machine at 20°C. Each experiment was repeated three times. Data analysis was performed on the Origin ITC200 software. Statistical analyses were performed with Prism (GraphPad Software). Comparisons were done by one-way ANOVA, followed by Dunnett’s multiple comparison tests with 99 % confidence interval. An adjusted *P* value <0.01 was considered statistically significant (** *P*<0.01, *** *P*<0.001, **** *P*<0.0001).

For measuring the affinity of the PilA-PilC interaction, the two purified proteins were buffer-exchanged into the same buffer (20 mM HEPES pH 7.5, 150 mM NaCl). The sample cell was loaded with 20 µM PilC, while the syringe was filled with 200 µM PilA. Each ITC run consisted of 20 titrations in total – the first injection was 0.4 µl, while all subsequent injections were 1.9 µl in volume. There was a 200-sec-long gap between each titration. To quantify the protein-sugar interactions, the purified PilC_Δpilin_ and the 3’-SL and 3’-SLN sugar ligands (Dextra Laboratories) were prepared in the same buffer (20 mM HEPES pH 7.6, 50 mM NaCl). The sample cell was loaded with 0.1 mM purified protein, and the syringe was filled with 2 mM of glycan ligand. Each ITC run consisted of 30 titrations in total with a 150-sec-long gap between each titration. The first titration was 0.4 µl, while all subsequent titrations were 1.2 µl in volume.

### NMR assignment and chemical shift perturbations

A sample containing ^13^C, ^15^N labelled PilA at 1 mM in NMR buffer (10 mM Na_2_HPO_4_/NaH_2_PO_4_ pH 7, 150 mM NaCl, 5 % D_2_O) was used for the TROSY-based assignment experiments: HNCA, HNCOCA, HNCO, HNCACO, HNCACB, and CBCACONH. All data were collected at 25°C on a Bruker Avance III HD 800MHz triple resonance spectrometer with a cryoprobe. Experiments were processed using MddNMR^60^ for reconstruction after Non-Uniform Sampling, and NMRPipe^61^. Peak picking and assignments were performed in SPARKY^62^.

Chemical shift perturbation experiments were performed using TROSY-based HSQC experiments with samples containing either ^15^N labelled PilA at 1 mM (in NMR buffer) or ^15^N labelled PilA at 1 mM and unlabelled PilC at 0.5 mM (in NMR buffer). Chemical shift perturbations were determined using the equation:

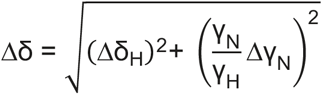

### Protein stability assays

To perform long-term protein stability tests, 6His-PilA and 6His-PilC were purified and mixed at 400 µM concentration. The single proteins and the complex – three independent aliquots for each – were kept at 4°C for four weeks. Samples were taken once a week, mixed with 2x Laemmli buffer and analysed by SDS-PAGE/Coomassie staining to reveal protein degradation.

To perform trypsin sensitivity assays, 6His-PilA and 6His-PilC were purified and one hundred µl aliquots of PilA, PilC and the PilA-PilC complex were prepared at 80 µM concentration. The PilA-PilC complex was incubated at 4°C O/N. The next day, the three aliquots were incubated with trypsin (Sigma) at 1/1,000 dilution for 60 min on ice. Samples of 10 µl were taken at 1, 5, 10, 20, 40 and 60 min. The samples were immediately mixed with 2x Laemmli buffer, boiled at 100°C and analysed by SDS-PAGE/Coomassie staining. The trypsin sensitivity assays were performed three times with freshly purified proteins.

### Glycan microarrays

The binding specificities of 6His-PilC, 6His-PilC_Δpilin_ and 6His-PilC^SK36^ – purified and concentrated to 1 mg/ml in 10 mM HEPES pH 7.5, 150 mM NaCl, 5 mM CaCl_2_ – were analysed using a NGL-based microarray system^42^. A previously described broad-spectrum microarray of 672 sequence-defined lipid-linked glycan probes was used^63^. The list of glycan probes is given in the Supplementary dataset 1. Details of the preparation of the glycan probes and the generation of the microarrays are listed in Table S3 in accordance with the MIRAGE guidelines (Minimum Information Required for A Glycomics Experiment)^64^. The microarray analyses were performed essentially as described^65^. In brief, after blocking of the slides for 1 h with HBS buffer (10 mM HEPES pH 7.4, 150 mM NaCl) containing 1 % (w/v) BSA (Sigma), 0.02 % (w/v) Casein (Pierce), and 10 mM CaCl2, the 6His-tagged PilC proteins were analysed under two conditions. In condition A, the microarrays were overlaid with the 6His-PilC proteins for 90 min as precomplexed protein-antibody complexes. These were prepared by preincubating the His-tagged PilC with mouse monoclonal anti-poly-histidine and biotinylated anti-mouse IgG antibodies (both from Sigma) at a ratio of 1:1.5:1.5 (by weight) and diluted in the blocking solution to provide a final PilC concentration of 50 µg/ml. In condition B, the microarrays were first overlaid with the 6His-PilC proteins at 100 µg/ml. This was followed by incubation with mouse monoclonal anti-His and biotinylated anti-mouse IgG antibodies (both at 10 µg/ml). In both conditions, binding was detected with Alexa Fluor-647-labelled streptavidin (Molecular Probes) at 1 µg/ml for 30 min. All steps were carried out at ambient temperature except for the precomplexation step which was carried out on ice. Imaging and data analysis are described in Table S3.

### Bioinformatics and modelling

Protein sequences were routinely analysed using DNA Strider^66^. Prediction of protein domains was done by interrogating the InterPro database with InterProScan^67^. Molecular visualisation of 3D structures was done using PyMOL (Schrödinger), which was used for generating the figures in this manuscript. The DALI server was used for comparing protein structures in 3D^68^. Protein 3D structures were downloaded from the RCSB PDB server. The 3d-SS^69^ server was used to superpose 3D protein structures with the STAMP algorithm^70^. PDBePISA^71^ was used for the exploration of macromolecular interfaces. Modelling was done using AlphaFold^38^ and AlphaFold-Multimer^72^ on Google Colab with default parameters.

## Acknowledgements

This work was funded by the Medical Research Council (MR/P022197/1). The glycan microarray studies were performed in the Carbohydrate Microarray Facility at the Glycosciences Laboratory, which is supported by Wellcome Trust biomedical resource grants (099197/Z/12/Z, 108430/Z/15/Z, and 218304/Z/19/Z) and partially by the March of Dimes Prematurity research centre grant (22-FY18-82). The sequence-defined glycan microarrays contain many saccharides provided by collaborators whom we thank, as well as members of the Glycosciences Laboratory for their contribution in the establishment of the NGL-based microarray system. We acknowledge the use of the crystallisation facility at Imperial College London, which is supported by the Biotechnology and Biological Sciences Research Council (BB/D524840/1) and Wellcome Trust (202926/Z/16/Z). We thank Angelika Gründling (Imperial College London), Sophie Helaine (Harvard Medical School) and Romé Voulhoux (CNRS Marseille) for critical reading of the manuscript.

