## Supplemental information for "Full structure/function analysis of all the pilin subunits in a type 4 pilus: a complex of minor pilins in *Streptococcus sanguinis* mediates binding to glycans"

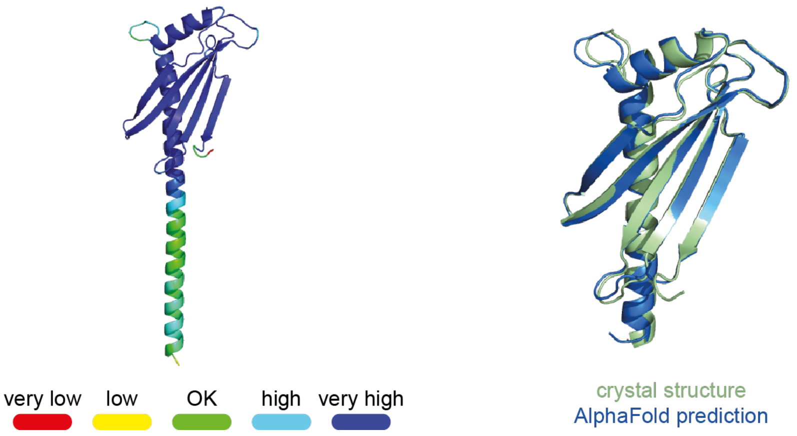


**Fig. S1. AlphaFold model of PilA. Left**, full-length model coloured following B-factor accuracy. **Right**, superposition of our crystal structure (green) with the corresponding portion of the AlphaFold[1] prediction (blue). The two structures superpose with an RMSD of 0.58 Å, showing structural identity.

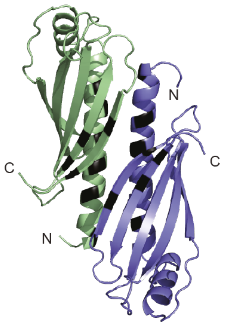


| H-bond | PilA 1 | dist. (Å) | PilA 2 |
|---|---|---|---|
| 1 | TRP$_{60}$ | 3.49 | THR$_{39}$ |
| 2 | HIS$_{141}$ | 2.83 | TYR$_{45}$ |
| 3 | TYR$_{50}$ | 3.55 | THR$_{53}$ |
| 4 | TYR$_{50}$ | 2.62 | GLN$_{57}$ |
| 5 | LYS$_{49}$ | 2.68 | TYR$_{139}$ |
| 6 | TYR$_{45}$ | 2.49 | GLU$_{155}$ |
| 7 | LEU$_{159}$ | 2.88 | GLU$_{155}$ |
| 8 | THR$_{39}$ | 3.38 | TRP$_{70}$ |
| 9 | TYR$_{45}$ | 2.84 | HIS$_{141}$ |
| 10 | TYR$_{139}$ | 2.85 | LYS$_{49}$ |
| 11 | GLU$_{155}$ | 2.56 | TYR$_{45}$ |
| 12 | GLU$_{155}$ | 2.88 | LEU$_{159}$ |

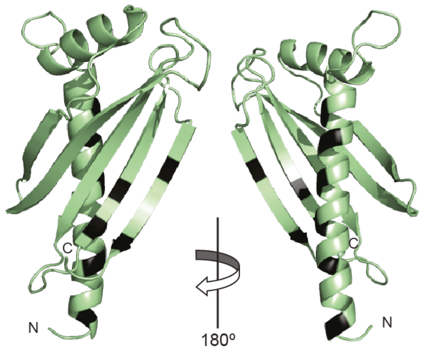


**Fig. S2. Interaction interface in PilA dimers. Left**, head-to-toe dimers in the crystal are stabilised by a series of hydrogen bonds between residues in α1-helix and the last 2 β-strands (black). Numbering is according to the full-length protein. **Right**, Close-up views at 180º rotation of one of the monomers in the dimer.

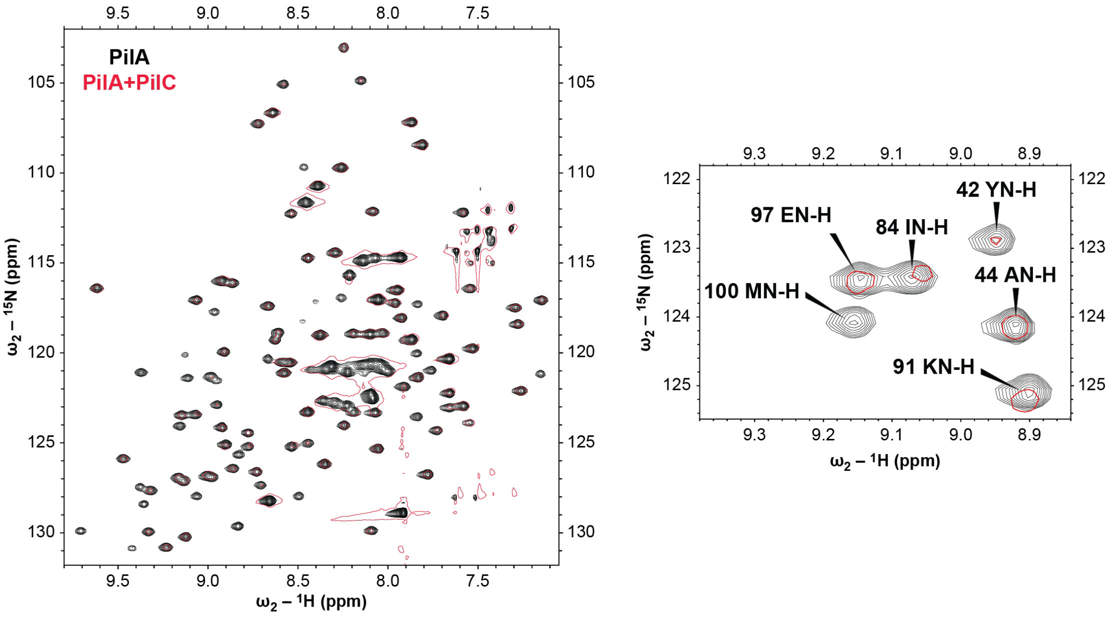


**Fig. S3. NMR analysis of PilA ± PilC. Left**, region of the $^{1}H$-$^{15}N$ HSQC spectrum of PilA. Free PilA spectrum is in black. A simplified PilA spectrum in the presence of PilC is in red. **Right**, close-up in an area of PilA showing significant amide shift changes in the presence of PilC. Numbering is according to the full-length protein.

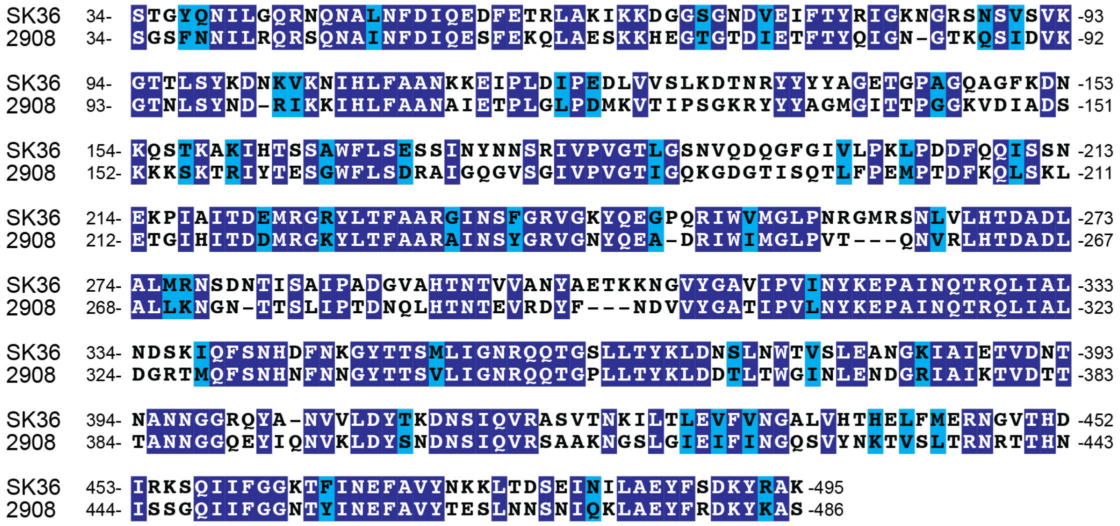


**Fig. S4. Sequence alignment of PilC from SK36 and 2908 strains, which were structurally characterised in this study**. Residues were shaded in dark blue (identical), light blue (conserved), or unshaded (different).

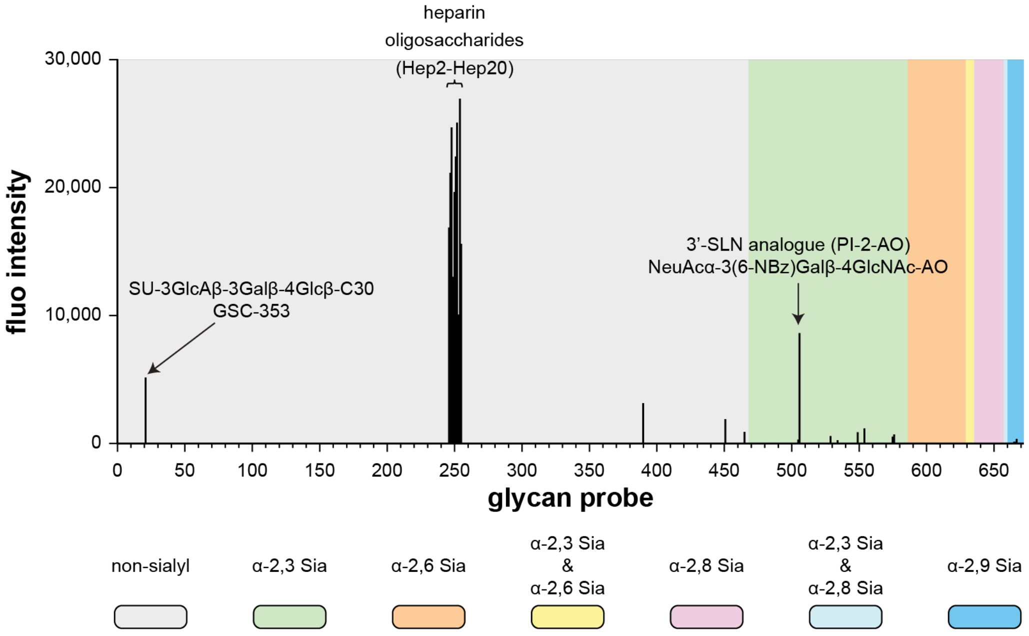


**Fig. S5. Glycan microarray analysis of the glycan-binding activity of PilC$_{\Delta pilin}$.** PilC$_{\Delta pilin}$ is the purified PilC protein without pilin module. The protein was analysed with and without precomplexation with detection antibodies, yielding similar results. The figure shows results with precomplexation. The results are the means of fluorescence intensities of duplicate spots, printed at 5 fmol/spot. In the glycan array the 672 lipid-linked probes are grouped according to sialyl linkages indicated by the coloured panels. The full list of glycan probes, their sequences, and binding scores are given in Supplementary dataset 1.

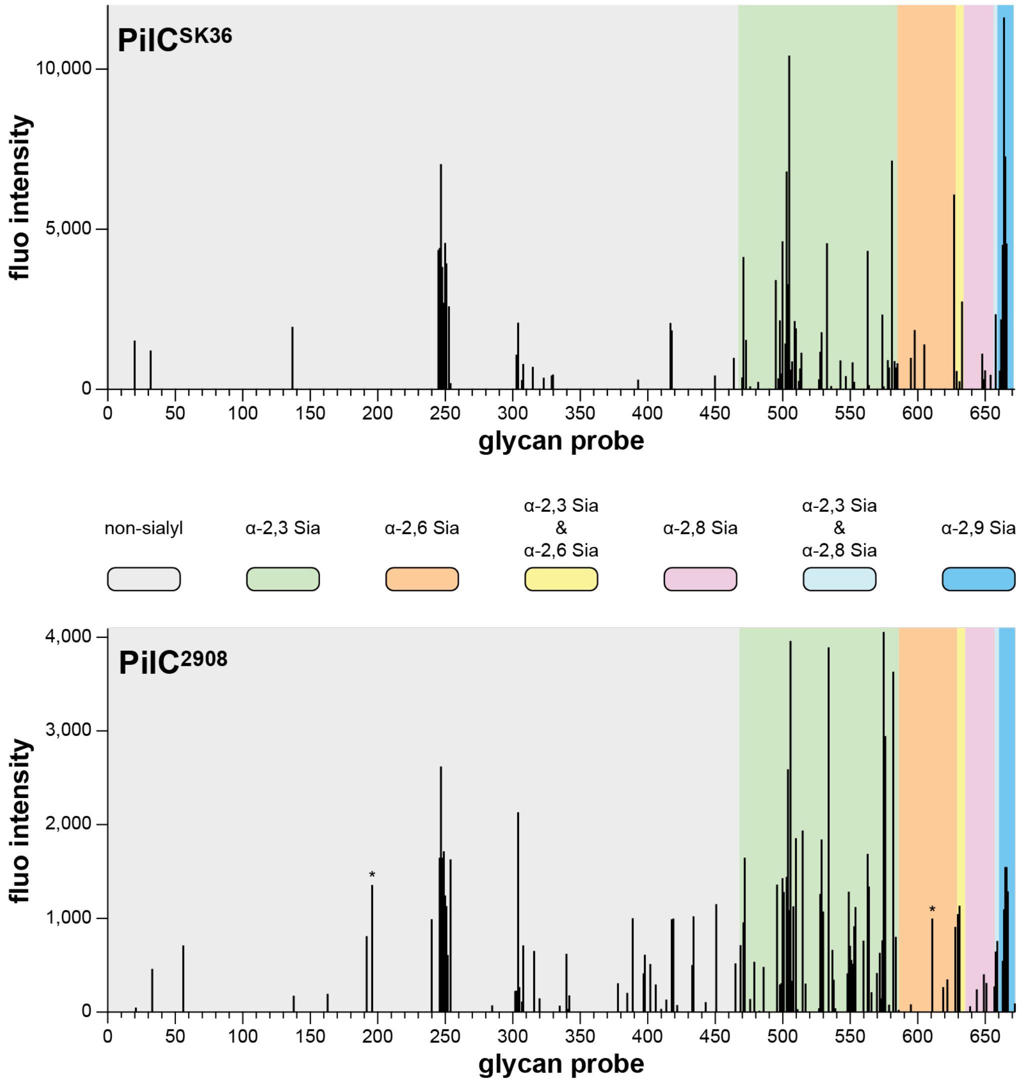


**Fig. S6. Comparison of the glycan microarray analyses of the glycan-binding activities of PilC$^{SK36}$ and PilC$^{2908}$ (full-length proteins).** The results are the means of fluorescence intensities of duplicate spots, printed at 5 fmol/spot. In the glycan array the 672 lipid-linked probes are grouped according to sialyl linkages as indicated by the coloured panels. The full list of glycan probes, their sequences, and binding scores are given in Supplementary dataset 1. For easy comparison of the sialyl glycan probes bound by the two proteins, a "condensed matrix" focused on sialyl glycan probes has been added in Supplementary dataset 1. * Signals with large error bars due to artefacts on the array slides.

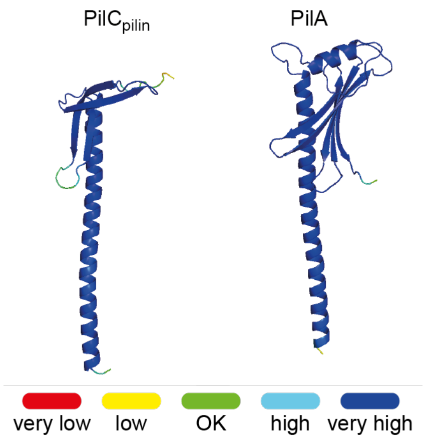


**Fig. S7. Accuracy of the two subunits in the AlphaFold model of the PilA-PilC$_{pilin}$ complex.** Predictions are coloured following B-factor accuracy.

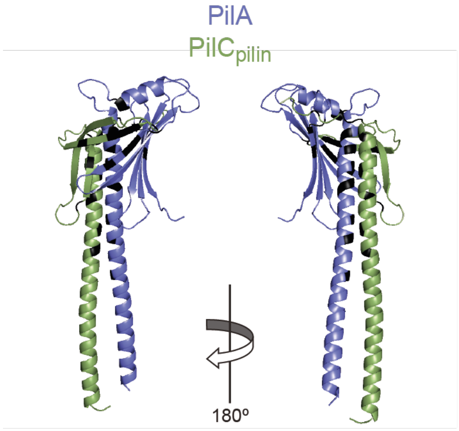


| H bond | PilC | dist. (Å) | PilA |
|---|---|---|---|
| 1 | GLN$_{43}$ | 3.05 | VAL$_{32}$ |
| 2 | LYS$_{65}$ | 2.67 | GLU$_{155}$ |
| 3 | TYR$_{79}$ | 2.16 | SER$_{73}$ |
| 4 | GLN$_{80}$ | 2.87 | LYS$_{93}$ |
| 5 | GLY$_{82}$ | 3.35 | PRO$_{91}$ |
| 6 | LYS$_{104}$ | 3.66 | TYR$_{43}$ |
| 7 | PHE$_{108}$ | 2.57 | SER$_{54}$ |
| 8 | ALA$_{110}$ | 3.72 | GLN$_{57}$ |
| 9 | GLY$_{118}$ | 3.04 | ILE$_{152}$ |
| 10 | ASN$_{47}$ | 3.65 | THR$_{39}$ |
| 11 | ILE$_{53}$ | 3.58 | TYR$_{50}$ |
| 12 | GLU$_{58}$ | 3.26 | LYS$_{49}$ |
| 13 | GLU$_{75}$ | 2.71 | HIS$_{96}$ |
| 14 | THR$_{78}$ | 2.48 | SER$_{94}$ |
| 15 | TYR$_{79}$ | 3.55 | VAL$_{74}$ |
| 16 | GLN$_{80}$ | 2.86 | LYS$_{93}$ |
| 17 | LYS$_{104}$ | 3.30 | TYR$_{43}$ |
| 18 | ILE$_{113}$ | 3.80 | TRP$_{60}$ |
| 19 | GLU$_{114}$ | 3.18 | GLN$_{57}$ |
| 20 | THR$_{115}$ | 3.34 | ILE$_{152}$ |
| 21 | GLY$_{118}$ | 3.60 | ARG$_{153}$ |

**Fig. S8. AlphaFold model of the PilA-PilC$_{pilin}$ complex. Left**, 180º view of the complex between PilA (blue) and the pilin module of PilC (green). Residues which stabilise the complex by forming a series of hydrogen bonds and salt bridges are highlighted in black. **Right**, the Table lists the residues forming H bonds, salt bridges are highlighted in red. Numbering is according to the full-length proteins.

**Table S1. Strains and plasmids used in this study.**

| Name | Details | Source |
|---|---|---|
| ***E. coli* strains** | | |
| DH5α | used for cloning | |
| BL21(DE3) | used for protein expression/purification | |
| B834(DE3) | used for SeMet protein expression/purification | |
| **Plasmids (all pET-28b derivatives)** | | |
| pET-28b | T7-based expression vector | Novagen |
| pET28-*pilA* | expressing 6His-PilA$_{33-164}$ | 2 |
| pET28-Strep-*pilA* | expressing Strep-PilA$_{33-164}$ | this study |
| pET28-Strep-*pilA*$_{I76S}$ | expressing Strep-PilA$_{33-164}$ with I76S mutation | this study |
| pET28-Strep-*pilA*$_{K93A}$ | expressing Strep-PilA$_{33-164}$ with K93A mutation | this study |
| pET28-Strep-*pilA*$_{T95A}$ | expressing Strep-PilA$_{33-164}$ with T95A mutation | this study |
| pET28-*pilB* | expressing 6His-PilB$_{36-461}$ | 2 |
| pET28-Strep-*pilB* | expressing Strep-PilB$_{36-461}$ | this study |
| pET28-*pilC* | expressing 6His-PilC$_{34-486}$ | 2 |
| pET28-*pilC*$^{SK36}$ | expressing 6His-PilC$_{34-495}$ from strain SK36 | this study |
| pET28-*pilC*$_{\Delta pilin}$ | expressing 6His-PilC$_{121-486}$ | this study |
| pET28-*pilC*$_{T356A}$ | expressing 6His-PilC$_{121-486}$ with T356A mutation | this study |
| pET28-*pilC*$_{K358A}$ | expressing 6His-PilC$_{121-486}$ with K358A mutation | this study |
| pET28-*pilC*$_{T364A}$ | expressing 6His-PilC$_{121-486}$ with T364A mutation | this study |
| pET28-*pilC*$_{E370A}$ | expressing 6His-PilC$_{121-486}$ with E370A mutation | this study |
| pET28-*pilC*$_{R374A}$ | expressing 6His-PilC$_{121-486}$ with R374A mutation | this study |
| pET28-*pilC*$_{N386A}$ | expressing 6His-PilC$_{121-486}$ with N386A mutation | this study |
| pET28-*pilC*$_{G389S}$ | expressing 6His-PilC$_{121-486}$ with G389S mutation | this study |
| pET28-*pilC*$_{Q390A}$ | expressing 6His-PilC$_{121-486}$ with Q390A mutation | this study |
| pET28-*pilC*$_{Y392A}$ | expressing 6His-PilC$_{121-486}$ with Y392A mutation | this study |
| pET28-Strep-*pilC* | expressing Strep-PilC$_{34-486}$ | this study |
| pET28-*pilE1* | expressing 6His-PilE1$_{46-157}$ | 2 |
| pET28-Strep-*pilE1* | expressing Strep-PilE1$_{46-157}$ | this study |
| pET28-*pilE2* | expressing 6His-PilE2$_{46-150}$ | 2 |
| pET28-Strep-*pilE2* | expressing Strep-PilE2$_{46-150}$ | this study |

*pilB*, *pilC*, and *pilC*$^{SK36}$ are codon-optimised synthetic genes.

**Table S2. Primers used in this study.**

| Name | Sequence |
|---|---|
| *pilA*-pETF | ggg**ccatgg**atcatcatcatcatcatcaTGATACAGGGCAAAGCCAGAC |
| Strep-*pilA*-pETF | ggg**ccatgg**attggagccacccgcagttcgaaaagGATACAGGGCAAAGCCAGAC |
| *pilA*-pETR | ccc**gtcgac**TTACTTCTGTGCCGATCTCAA |
| *pilA*$_{A76S}$#1 | GGCAATCCATCATCGGTTTATTCAGAGTTTGACGAGCGGGCCC |
| *pilA*$_{A76S}$#2 | GGGCCCGCTCGTCAAACTCTGAATAAACCGATGATGGATTGCC |
| *pilA*$_{K93A}$#1 | GATCCTTCAACAGAGCCGATTGCGTCAACCCATACCTTCAAAG |
| *pilA*$_{K93A}$#2 | CTTTGAAGGTATGGGTTGACGCAATCGGCTCTGTTGAAGGATC |
| *pilA*$_{T95A}$#1 | CAACAGAGCCGATTAAGTCAGCCCATACCTTCAAAGATGGC |
| *pilA*$_{T95A}$#2 | GCCATCTTTGAAGGTATGGGCTGACTTAATCGGCTCTGTTG |
| *pilB*-pETF | ggg**ccatgg**atcatcatcatcatcatcaTAGCAGCCGTGAACTGATTGA |
| Strep-*pilB*-pETF | ggg**ccatgg**attggagccacccgcagttcgaaaagAGCAGCCGTGAACTGATTGA |
| *pilB*-pETR | ccc**ggatcc**TTACGGACCGCTAACAAACC |
| *pilC*-pETFbis | ggg**ccatgg**atcatcatcatcatcatcaTAGCGGCAGCTTTAATAACATTCTGCG |
| Strep-*pilC*-pETF | ggg**ccatgg**attggagccacccgcagttcgaaaagAGCGGCAGCTTTAATAACATTCTGCG |
| *pilC*-pETR | ccc**ggatcc**TTAGCTTGCTTTGTATTTATCGC |
| *pilC*$_{\Delta pilin}$ | ggg**ccatgg**atcatcatcatcatcatcaTGATATGAAAGTGACCATTCCGAGCGGTAAACGTTATTAC |
| *pilC*$_{T356A}$#1 | CAGCAGACAGGTCCGCTGCTGGCATATAAACTGGATGATACCC |
| *pilC*$_{T356A}$#2 | GGGTATCATCCAGTTTATATGCCAGCAGCGGACCTGTCTGCTG |
| *pilC*$_{K358A}$#1 | GGTCCGCTGCTGACATATGCACTGGATGATACCCTGAC |
| *pilC*$_{K358A}$#2 | GTCAGGGTATCATCCAGTGCATATGTCAGCAGCGGACC |
| *pilC*$_{T364A}$#1 | CTGGATGATACCCTGGCCTGGGGTATTAATC |
| *pilC*$_{T364A}$#2 | GATTAATACCCCAGGCCAGGGTATCATCCAG |
| *pilC*$_{N368A}$#1 | CCCTGACCTGGGGTATTGCTCTGGAAAATGATGGTC |
| *pilC*$_{N368A}$#2 | GACCATCATTTTCCAGAGCAATACCCCAGGTCAGGG |
| *pilC*$_{E370A}$#1 | CCTGGGGTATTAATCTGGCAAATGATGGTCGCATTGC |
| *pilC*$_{E370A}$#2 | GCAATGCGACCATCATTTGCCAGATTAATACCCCAGG |
| *pilC*$_{R374A}$#1 | GGTATTAATCTGGAAAATGATGGTGCCATTGCCATCAAAACCGTTGATAC |
| *pilC*$_{R374A}$#2 | GTATCAACGGTTTTGATGGCAATGGCACCATCATTTTCCAGATTAATACC |
| *pilC*$_{G389S}$#1 | CCACCACCGCAAATAATGGTTCTCAAGAATATATCCAGAACG |
| *pilC*$_{G389S}$#2 | CGTTCTGGATATATTCTTGAGAACCATTATTTGCGGTGGTGG |
| *pilC*$_{Q390A}$#1 | CACCGCAAATAATGGTGGTGCAGAATATATCCAGAACGTG |

| | |
|---|---|
| *pilC*$_{Q390A}$#2 | CACGTTCTGGATATATTCTGCACCACCATTATTTGCGGTG |
| *pilC*$_{Y392A}$#1 | CAAATAATGGTGGTCAAGAAGCTATCCAGAACGTGAAACTGG |
| *pilC*$_{Y392A}$#2 | CCAGTTTCACGTTCTGGATAGCTTCTTGACCACCATTATTTG |
| *pilC*$^{SK36}$-pETF | ggg**ccatgg**atcatcatcatcatcatcatAGCACCGGTTATCAGAACATTCTG |
| *pilC*$^{SK36}$-pETR | ccc**ggatcc**TTATTTGGCACGGTATTTATCGC |
| *pilE*-pETF | gg**ccatgg**atcatcatcatcatcatcatcaAGATAACGCTCGTAAGAGCC |
| Strep-*pilE*-pETF | ggg**ccatgg**attggagccacccgcagttcgaaaagcaAGATAACGCTCGTAAGAGCC |
| *pilE1*-pETR | cc**ggatcc**TTAGTTTGAGTTTACACCATTAGCAGA |
| *pilE2*-pETR | cc**ggatcc**TTATTTTGAATTAGCACCAGCTTCG |

*pilB*, *pilC*, and *pilC*$^{SK36}$ are codon-optimised synthetic genes. Overhangs are in lower case. Restriction sites are in bold. Mismatches are in red.

**Table S3. Glycan microarray specifications following the MIRAGE guidelines[3].**

| | |
|---|---|
| **1. Glycan binding sample** | |
| Description of sample | *Sample names*: PilC, $PilC_{\Delta pilin}$ and $PilC^{SK36}$. *Origin*: recombinant. All samples contained an N-terminal 6His tag. *Method of preparation*: see "Glycan microarray" section in the main text. |
| Sample modifications | Not relevant. |
| Assay protocol | Microarray analyses were performed essentially as described[4], for modifications of the protocol see "Glycan microarray" section in the main text. |
| **2. Glycan library** | |
| Glycan description for defined glycans | A broad-spectrum screening microarray containing 672 sequence-defined oligosaccharide probes was used. The probe names, corresponding sequences, and IDs in the international glycan structure repository GlyTouCan (https://glytoucan.org/) are displayed in Supplementary dataset 1. These probes represent a subset of a recently generated large screening microarray containing around 900 glycan probes (in-house designation "Array sets 42-56", which will be published elsewhere). The NGL probes are from the collection assembled by the Glycosciences Laboratory (https://glycosciences.med.ic.ac.uk/glycanLibraryList.html). |
| Glycan description for undefined glycans | Not relevant. |
| Glycan modifications | No modification was carried out for natural glycolipids. NGLs, unless otherwise specified, were prepared by reducing oligosaccharides by reductive amination with the amino lipid, 1,2-dihexadecyl-*sn*-glycero-3-phosphoethanolamine (DHPE)[5]. AO indicates NGLs prepared by reducing oligosaccharides by oxime ligation with an aminooxy functionalised DHPE (AOPE)[6]. For full description on the definition of lipid moieties of the glycan probes see https://glycosciences.med.ic.ac.uk/docs/lipids.pdf. |
| **3. Printing surface** | |
| Description of surface | Nitrocellulose-coated glass microarray slides. |
| Manufacturer | 16-pad UniSart 3D Microarray Slide from Sartorius (Goettingen, Germany). |
| Custom preparation of surface | Not relevant. |
| Non-covalent Immobilisation | The lipid-linked oligosaccharide probes for arraying and non-covalent immobilisation on nitrocellulose-coated glass slides[4] were formulated as liposomes by adding carrier lipids, 1,2-dihexanoyl-*sn*-glycero-3-phosphocholine (DHPC) and cholesterol. |
| **4. Arrayer (Printer)** | |
| Description of arrayer | Nano-Plotter 2.1 from GeSiM (Radeberg, Germany). |

| | |
|---|---|
| Dispensing mechanism | Non-contact liquid delivery with four dispensing tips. |
| Glycan deposition | Approximately 0.33 nl was printed per spot. Lipid-linked glycan probes were printed at 2 and 5 fmol per spot in duplicate. |
| Printing conditions | The printing solutions were all aqueous based. Printing was performed at ambient temperature and relative humidity of 58 %. In addition to the lipid-linked glycan probes, the "liposome" printing solutions contained 100 pmol/µl of DHPC and cholesterol as lipid carriers (both from Sigma). The concentrations of the lipid-linked glycan probes were 5 and 15 pmol/µl for the 2 and 5 fmol per spot levels, respectively. The printing solutions also contained Cy3 NHS ester (GE Healthcare) at 20 ng/ml (26 fmol/µl) as a marker to monitor the printing process. |
| **5. Glycan microarray with "map"** | |
| Array layout | Each array slide contained 16-pad subarrays. Each pad was set up for printing 64 probes maximum, each at 2 levels in duplicate (four spots for one probe in a row); 256 spots (16x16) in total in each pad. The 672 lipid-linked probes in the screening arrays were printed on multiple subarrays for parallel binding analyses. |
| Glycan identification and quality control | The quality control of the screening microarrays of sequence-defined glycan probes was carried out with a panel of biotinylated plant lectins (Vector Laboratories), *e.g.*, *Ricinus communis* Agglutinin I, *Aleuria aurantia* lectin, Concanavalin A, and wheat germ agglutinin, a wide range of anti-carbohydrate antibodies, and several commercial bacterial adhesins and toxins. A number of viral adhesive proteins that have been published previously were also analysed, including VP1 proteins of polyomaviruses – simian virus 40[7], human JC polyomavirus[8] and BK polyomavirus[9] – human adenovirus 52 fibre knob[10], and Chikungunya virus[11]. These data will be described elsewhere and are available upon request. |
| **6. Detector and data processing** | |
| Scanning hardware | GenePix 4300A from Molecular Devices (UK). |
| Scanner settings | *Scanning resolution*: 10 µm/pixel.<br>*Laser channel*: Red (scan wavelength 635 nm).<br>*PMT*: 350.<br>*Scan power*: various laser powers were used (indicated in Supplementary dataset 1) to record optimal scan images without spot saturation. |
| Image analysis software | GenePix® Pro 7 (Molecular Devices). |
| Data processing | The gpr files were entered into an in-house microarray database using software designed by Mark Stoll for data processing (http://www.beilstein-institut.de/en/publications/proceedings/glyco-2009). No particular normalisation method or statistical analysis was used for the results of the screening arrays. |
| **7. Glycan microarray data presentation** | |
| Data presentation | The microarray binding results are presented as histogram charts in Fig. 7A, Fig. S5 and Fig. S6. The results table with binding scores and relative binding intensities shown as "matrix" are in Supplementary dataset 1. This dataset also displays a focused |

| | |
|---|---|
| | matrix of the bound sialyl glycan probes with their sequences. |
| **8. Interpretation and conclusion from microarray data** | |
| Data interpretation | No software or algorithms were used to interpret processed data. |
| Conclusions | PilC , $PilC_{\Delta pilin}$ and $PilC^{SK36}$ bound to sialyl glycans and sulphated GAGs (mainly heparin) NGL probes in the screening array. $PilC_{\Delta pilin}$ showed much stronger binding to heparin NGL probes relative to sialyl probes. For the full-length proteins, PilC and $PilC^{SK36}$, very similar binding patterns were observed to a broad range of sialylated glycans, mainly sialyl α2-3-linked and α2-9-linked polysialic. Among the differences there is the probe NeuAcα-6GalNAc-AO bound strongly by $PilC^{SK36}$ and weakly by PilC. |

**Supplementary dataset legend**

**Supplementary dataset 1. Results of the different glycan array experiments, with detailed list of probes on the array.** This includes experiments performed with PilC, PilC$_{\Delta pilin}$ and PilC$^{SK36}$.
